# Mechanisms of superior respiratory IgA responses against SARS-CoV-2 after mucosal vaccination

**DOI:** 10.1101/2025.09.22.677712

**Authors:** Jinyi Tang, Arka Sen Chaudhuri, Panke Qu, Pei Li, Yajie Liu, Yue Wu, Kristin Wavell, MarthaJoy M. Spano, In Su Cheon, Chaofan Li, Jane Yu, Harish Narasimhan, Mohd Arish, Wei Qian, Gislane de Almeida Santos, Samuel P. Young, Anika Agarwal, Fangming Zhu, Takao Kobayashi, Maria Velegraki, Mark H. Kaplan, Justin J. Taylor, Guizhi Zhu, Haitao Hu, Zihai Li, Hui Hu, Hirohito Kita, Nu Zhang, Shan-Lu Liu, W. Gerald Teague, Jie Sun

**Author notes:** Correspondence to: Jie Sun. Co-first authors (Jinyi Tang, Arka Sen Chaudhuri). Co-senior authors (Shan-Lu Liu, W. Gerald Teague, Jie Sun).

## Abstract

Mucosal immunization and respiratory IgA offer significant promise in protecting against airborne pathogens, including SARS-CoV-2. However, the conditions and mechanisms that lead to the robust induction of respiratory IgA responses following mucosal vaccination remain poorly understood. It is also currently debatable whether mucosal vaccination is still warranted given that most individuals in developed countries have established a hybrid immunity from vaccination and infection. Here we characterized respiratory mucosal immune responses after SARS-CoV-2 infection, vaccination or both in humans. We found that hybrid immunity resulted in moderately increased respiratory IgA and neutralizing antibody responses compared to infection or vaccination alone. However, a direct comparison of hybrid immunity and a mucosal adenovirus-based booster vaccination in animal models revealed that respiratory booster immunization elicited markedly stronger and more durable respiratory IgA, T cell response, and protective immunity against SARS-CoV-2, supporting the promise of respiratory mucosal vaccination. Mechanistically, we found that mucosal booster immunization induced local IgA- secreting cells in the respiratory mucosa, aided by pulmonary CD4^+^ T cells *in situ*. Strikingly, local IL-21-producing Blimp-1^+^ Th1 effector cells were critical in mediating the CD4^+^ T cell help for respiratory IgA production. Furthermore, lung macrophages were important for this respiratory IgA response via the production of TGF-β. Consequently, we demonstrated delivery of adenoviral booster to the lower airway was necessary to generate robust upper and lower airway IgA responses. Collectively, our results uncover a local cellular network supporting enhanced respiratory IgA responses, with implications for the development of optimal mucosal immunization strategies against SARS-CoV-2 and other respiratory pathogens.

## Introduction

The current intramuscular SARS-CoV-2 mRNA vaccines have demonstrated efficacy in eliciting strong immune responses in the circulation, significantly reducing the severity of disease following SARS-CoV-2 infection ^1^. Nevertheless, a key limitation is their relative inability to induce robust respiratory mucosal immunity ^2–4^. To overcome this, the Coronavirus Vaccines R&D roadmap and Project NextGen have launched efforts to develop next-generation SARS-CoV-2 mucosal vaccines ^5,6^. However, the lack of clear understanding of the approaches and mechanisms underlying the robust respiratory mucosal immunity poses significant hurdles for the development of efficacious mucosal vaccines against SARS-CoV-2 and other airborne pathogens.

It has been shown that SARS-CoV-2 infection in mRNA-vaccinated individuals provides greater protection against subsequent breakthrough infections compared to vaccination or infection alone ^7,8^, indicating that hybrid immunity confers broader and more durable protective responses against SARS-CoV-2 variants. Current evidence also suggests a similar trend for neutralizing antibody (nAb) responses, which are stronger and more durable in the case of vaccination plus infection ^9–12^. Nevertheless, the distinguishing characteristics of hybrid immunity in the respiratory tract, the primary site of infection, remain to be fully examined. Furthermore, the widespread prevalence of hybrid immunity raises questions about the continued necessity of a mucosal booster. It remains unclear whether such a booster would substantially enhance mucosal immunity and protection against virus variants compared to hybrid immunity.

In the context of SARS-CoV-2, evidence suggests that mucosal IgA is a more robust indicator of protective immunity against breakthrough infections, rather than circulating or mucosal IgG antibodies ^13–15^. However, there is a significant knowledge gap regarding the strategies and mechanisms eliciting effective respiratory mucosal IgA production after infection or vaccination. Recent data have suggested that respiratory IgA-secreting cells are programmed by follicular helper T (Tfh) cells in regional lymphoid tissues and migrate to the respiratory mucosa ^16^. It is currently unknown whether those migrating IgA-secreting cells require additional help at the mucosal surface, either from other mucosa-homing immune cells or periphery-homing terminal effector T cells. In this report, we have performed a comparative analysis of the protective mucosal immunity induced by hybrid immunity and/or a mucosal booster vaccination. We discovered that mucosal booster vaccination induces robust respiratory IgA and protective immunity. Additionally, we identified a local cellular network involving pulmonary Blimp-1-expressing CD4^+^ Th1 effector cells and lung macrophages which are critical for supporting mucosal IgA responses after respiratory immunization.

## Results

### Hybrid immunity is characterized by enhanced respiratory IgA and nAb

To determine the respiratory mucosal signature of hybrid immunity, we examined blood, bronchoalveolar lavage (BAL), and nasal wash specimens from a cohort of children with minor airway structural anomalies and treatment-recurrent recurrent wheeze who underwent clinically indicated bronchoscopy and were assigned to four categories according to COVID-19 and vaccination status. The categories included COVID-19 naïve, COVID-19 vaccinated with no evidence of infection, COVID-19 convalescents alone, and COVID-19 vaccinated individuals with SARS-CoV-2 infection history (hybrid) (Fig. 1a). Detailed cohort information is included in Extended Data tables 1 and 2. Since majority of the pediatric infection were mild or non- symptomatic ^17^, we performed infection history adjustment based on the levels of SARS-CoV-2 N-specific IgG by enzyme-linked immunoassay (ELISA) in the blood and BAL (Extended Data Fig. 1a). Subsequently, we compared SARS-CoV-2 S1 or receptor binding domain (RBD)- specific IgG and IgA levels in the plasma, BAL and nasal wash. As previously reported ^3,12,18,19^, SARS-CoV-2 infection, and particularly vaccination alone, induced robust S1 or RBD-specific plasma, BAL, and nasal IgG levels (Fig. 1b-d). Similarly, vaccination plus infection induced high levels of S1 or RBD-specific IgG systemically in the blood or in the respiratory mucosa, at least the same levels or higher than those of the infected or vaccinated group (Fig. 1b-d). The S1 or RBD-specific IgA levels in the plasma were elevated in the infection, vaccinated, or hybrid groups, with the hybrid group tending to have higher IgA levels (Fig. 1e). As we have previously reported ^3^, natural infection, but not mRNA vaccination, induced BAL IgA responses (Fig. 1f).

**Fig. 1:**
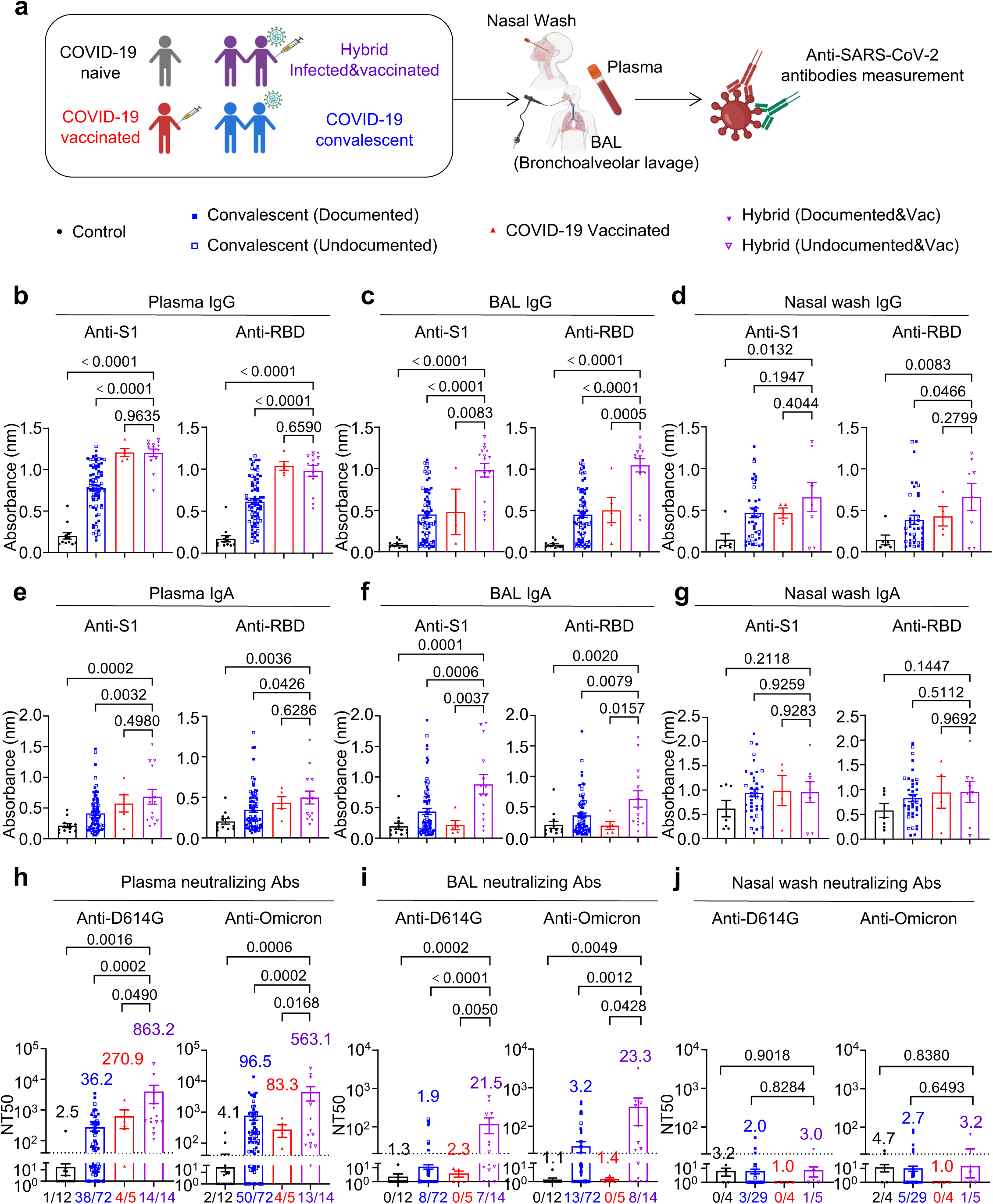
Systemic and respiratory mucosal Ab responses in individuals with SARS-CoV-2 vaccination and infection. **a,** Schematic of recruited cohorts and experimental procedures. Figures were created with BioRender. **b-g**, Levels of SARS-CoV-2 S1 or RBD binding IgG **(b-d)** or IgA **(e-g)** in plasma **(b,e)**, BAL **(c,f)**, and nasal wash **(d,g)** of COVID-19 naïve individuals (n = 12), COVID-19 convalescents (n = 75), COVID-19 vaccinated individuals (n = 5), vaccinated plus infected individuals (n = 14). **h-j**, Levels of SARS-CoV-2 neutralizing Ab against D614G and Omicron BA.1 strains in plasma **(h)**, BAL **(i)**, and nasal wash **(j)** of COVID-19 naïve individuals (n = 14 for plasma, n = 12 for BAL and n = 4 for nasal wash), COVID-19 convalescents (n = 72 for plasma and BAL, n = 29 for nasal wash), vaccinated individuals (plasma and BAL, n = 5; nasal wash, n = 3), or vaccinated plus infected individuals (plasma and BAL, n = 14; nasal wash, n =5). Individuals with undocumented COVID-19 infection are indicated as hollow symbols in groups of COVID-19 convalescents and COVID-19 vaccinated plus infected. Enrolled donors’ demographics are provided in Extended table 1. **b-j**, One-way ANOVA with multiple- comparisons test. Data presented as mean ± s.e.m.

BAL IgA levels in the hybrid group were particularly higher than those in the infection or vaccinated individuals (Fig. 1f). In contrast, no significant IgA response was observed in the nasal washes (Fig. 1g). Of note, respiratory IgG and IgA levels, except nasal IgA levels, were generally correlated with blood IgG and IgA levels (Extended Data Fig. 1b-g).

We next examined the plasma, BAL, and nasal neutralizing Ab activity against SARS-CoV-2 D614G and Omicron BA.1 spike-pseudotyped lentiviruses ^20^ (Fig. 1h-j). Since most COVID-19 convalescent individuals were infected during the Omicron wave, plasma neutralization titer (NT_50_) in those individuals was generally higher against Omicron BA.1 than the ancestral D614G (Fig. 1h). In contrast, vaccinated individuals had higher neutralizing activity against D614G compared to the Omicron variant (Fig. 1h). Notably, the hybrid group had increased neutralizing Ab levels against both D614G and Omicron in the plasma compared to those in the infection or vaccination groups (Fig. 1h). In the BAL and nasal washes, vaccination alone failed to induce any significant levels of neutralizing Abs against D614G or Omicron, while prior infection elicited minimal if any levels of neutralizing Abs (Fig. 1i, j). However, the hybrid group induced clearly increased levels of neutralizing Abs against D614G and Omicron compared to vaccination or infection alone in the BAL, but not in the nasal wash (Fig. 1i, j). Together, these data indicate that hybrid immunity can elicit IgA responses and neutralizing immunity in the lower respiratory mucosal surface compared to mRNA vaccination and natural infection.

### Mucosal humoral, cellular and molecular signatures of hybrid immunity in animal models

To provide a mechanism for the development of protective mucosal immunity after SARS-CoV-2 vaccination and infection, we established mouse models of hybrid immunity (Fig. 2a). To achieve this, we immunized WT young C57BL/6 mice with 2 x Spike (S) mRNA/LNP (lipid nanoparticle) (mRNA-S) and subsequently challenged them with SARS-CoV-2 mouse adapted MA10 strain. mRNA-S immunization and MA10 infection alone served as vaccination-only and infection-only controls, respectively. Single-cell RNA sequencing (scRNA-seq) of BAL fluid revealed that hybrid immunity was associated with a markedly increased abundance of adaptive immune cells compared to innate myeloid cells (Fig. 2b, c and Extended Data Fig. 2a-c).

**Fig. 2:**
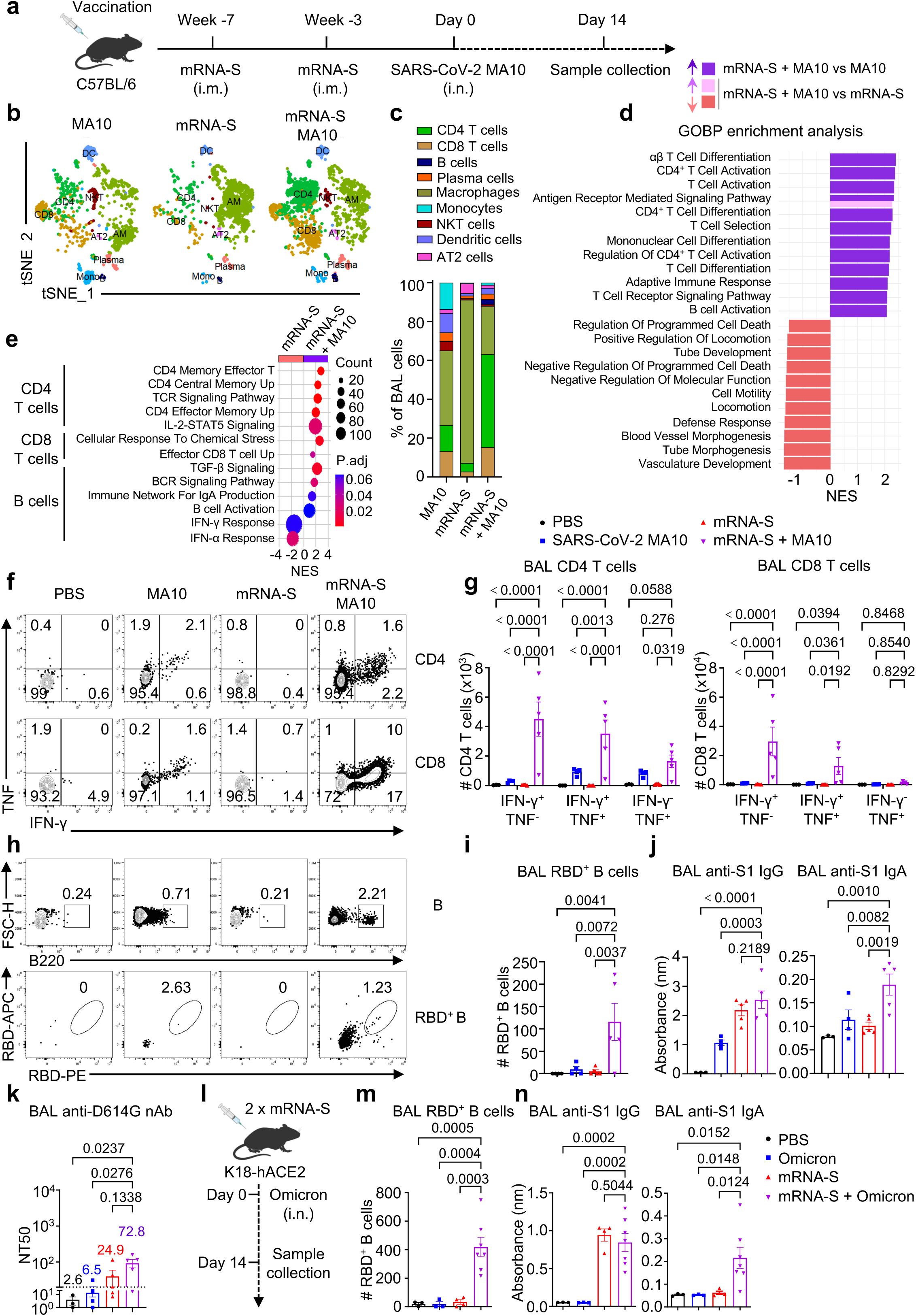
Mucosal signatures of hybrid immunity in animal models. **a,** Schematic of experimental design. C57BL/6 mice were immunized as indicated (PBS, n = 3; SARS-CoV-2 MA10, n = 4; mRNA-S, n = 5; mRNA-S + MA10, n =5). **b,c**, An scRNA-seq UMAP plot **(b)** with percentage of different clusters **(c)** of BAL cells from mice at day 6 post SARS- CoV-2 MA10 infection or without infection. **d**, GOBP enrichment pathways of BAL cells from scRNA-seq analysis. **e**, Enrichment of upregulated pathways in BAL CD4^+^, CD8^+^ T and B cells from mRNA-S+ MA10 group versus mRNA-S group. **f-i**, Represented flow cytometry plot **(f,h)** and summary **(g,i)** of SARS-CoV-2 peptide-specific CD4^+^, CD8^+^ T cell **(f,g)** and RBD-specific B cell responses **(h,i)** in the BAL from different groups. **j**, SARS-CoV-2 S1-specific IgG and IgA responses in the BAL from different groups. **k**, SARS-CoV-2-specific neutralizing Ab response against D614G in the BAL from different groups. **l**, Schematic of experimental design. *K18- hACE2* mice were immunized as indicated (PBS, n = 3; SARS-CoV-2 Omicron BA.1, n = 3; mRNA-S, n = 4; mRNA-S + Omicron BA.1, n =7). **m**, Flow cytometry summary of SARS-CoV-2 RBD-specific B cell responses in the BAL from different groups. **n**, SARS-CoV-2 S1-specific IgG and IgA responses in the BAL from different groups. **g**, Two-way ANOVA with multiple- comparisons test; **i-k,m,n**, one-way ANOVA with multiple-comparisons test. Data presented as mean ± s.e.m. **f-k,m,n**, Data are representative of two independent experiments with similar results.

Differential gene expression (DEG) and pathway analysis showed enrichment of processes associated with T- and B cell activation and host defense in the hybrid immunity group (Fig. 2d and Extended Data Fig. 2 d, e). Particularly, compared to those from mice immunized with mRNA-S alone, BAL T cells from the hybrid immunity group exhibited enrichment in gene sets associated with effector T cell differentiation and TCR signaling, while BAL B cells from the same group showed enhanced enrichment in pathways related to B cell activation, IgA production, and other related processes (Fig. 2e).

Consistent with the scRNA seq data, flow cytometry analysis demonstrated that hybrid immunity is associated with increased SARS-CoV-2 epitope-specific IFN-γ and TNF-producing effector CD4^+^ and CD8^+^ T cells at the respiratory mucosal surface (Fig. 2 f, g and Extended Data Fig. 3 a-c). Vaccination combined with MA10 infection also resulted in significantly elevated mucosal RBD-specific B cell levels (Fig. 2h, i and Extended Data Fig. 3 d, e). With regards to systemic and mucosal Ab responses, the hybrid immunity group exhibited higher mucosal S1- or RBD- specific IgG levels compared to the infection-alone group, however, responses were comparable to the vaccination-alone group (Fig. 2j and Extended Data Fig. 3f-g). Additionally, vaccination plus infection induced higher levels of IgA response compared to either infection or vaccination alone, particularly in the BAL (Fig. 2j and Extended Data Fig. 3h). As a result, the hybrid immunity generated the highest respiratory neutralizing Ab levels among the different groups examined (Fig. 2k).

We also established a hybrid immunity model using K18-hACE2 transgenic mice, allowing us to infect the mice with a clinical isolate of the Omicron strain (Fig 2l). Omicron infection in this model induced minimal lung pathology and immune responses ^21,22^. Nevertheless, we observed increased respiratory cellular (T and B cells) and elevated mucosal and plasma IgA levels, but similar RBD and S1-specific IgG antibodies, following vaccination plus Omicron infection, compared to those of vaccination or Omicron infection alone (Fig. 2m, n and Extended Data Fig. 4). Together these data revealed that, similar to the human study described above, hybrid immunity is also associated with increased mucosal adaptive cellular and humoral immune responses in mice compared to mRNA vaccination or infection alone.

### Comparison of respiratory protective immune responses between hybrid immunity and mucosal booster vaccination

The aforementioned human and mouse data suggest that hybrid immunity is capable of generating stronger respiratory mucosal immunity than the current vaccines. However, the prevalence of hybrid immunity in the real world raises a critical question on whether mucosal vaccination (likely a booster vaccination considering prior immunization via vaccine or infection) is still necessary. Thus, it is important to perform a side-by-side comparison of the protective mucosal immunity induced by hybrid immunity versus a mucosal booster vaccination. To this end, we compared mucosal humoral and cellular immune responses between mRNA vaccination plus MA10 infection and mRNA vaccination plus a replication-defective adenoviral construct expressing the ancestral Spike protein (Ad5-S) (Fig. 3a). Ad5-S mucosal booster immunization induced a marked enhancement in the percentage and quantity of IFN-γ producing Th1 cells, and IFN-γ or TNF-producing CD8^+^ T cells (Fig. 3b, c and Extended Data Fig. 5 a, b). Ad5-S mucosal booster also elicited greatly increased numbers of CD4^+^ and CD8^+^ tissue-resident memory T (T_RM_) cells (CD69^+^CD49a^+^ for CD4^+^ T cells or CD69^+^CD103^+^ for CD8^+^ T cells) (Extended Data Fig. 5 c, d), a cell type that mediates significant protection against respiratory viral infection ^23^. Furthermore, Ad5-S mucosal booster stimulated an increase in RBD-specific B cells, particularly RBD-specific IgA^+^ B cells and plasma cells, which lasted for at least two months post-booster immunization (Fig. 3d-f and Extended Data Fig. 5e).

**Fig. 3:**
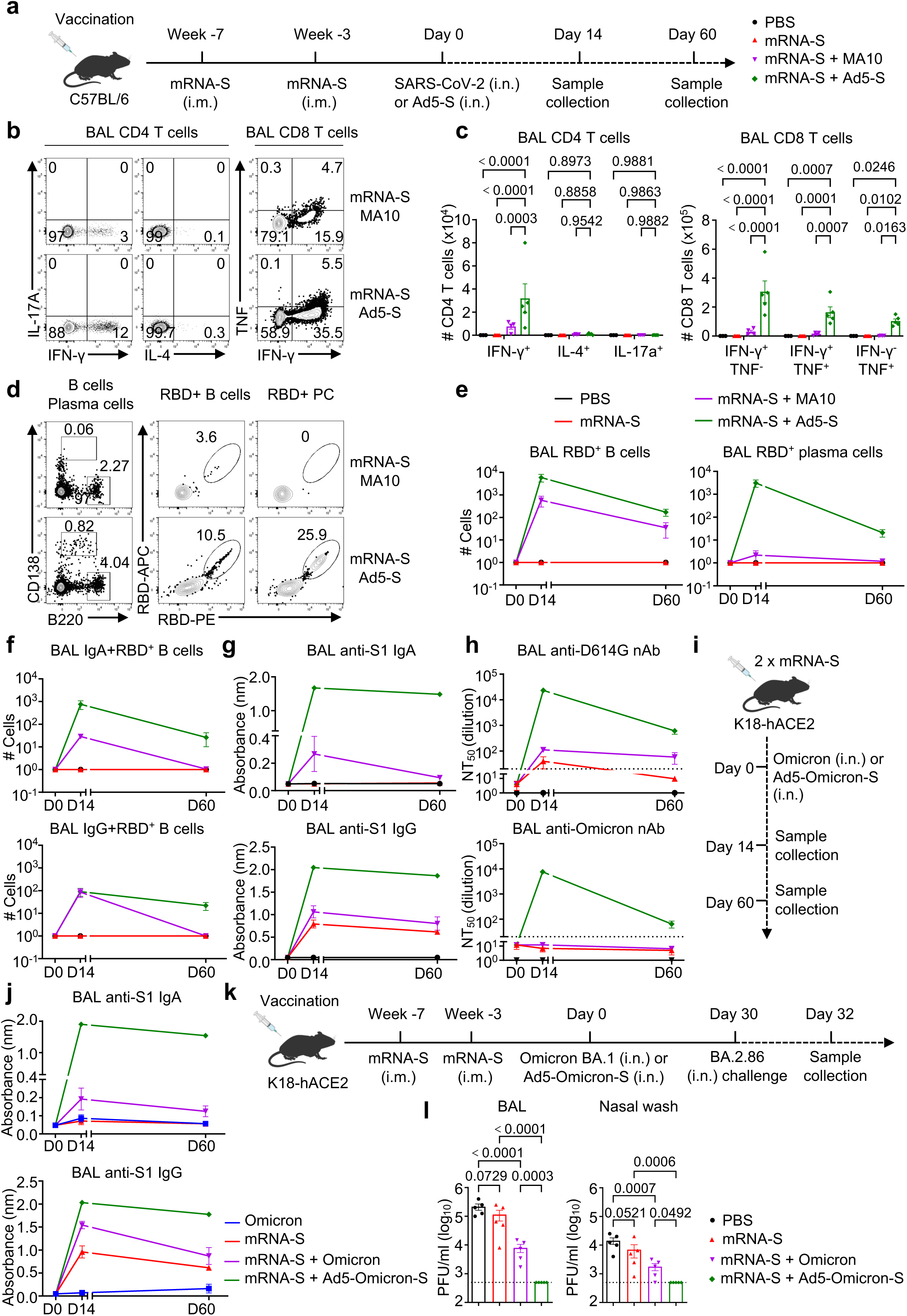
Comparative analyses of the mucosal protective responses after hybrid immunity and mucosal booster vaccination. **a,** Schematic of experimental design. C57BL/6 mice were immunized as indicated (Day 0, n = 3; Day 14, PBS, n = 3; mRNA-S, n = 5; mRNA-S + MA10, n =4; mRNA-S + Ad5-S, n =5; Day 60, PBS, n = 3; mRNA-S, n = 3; mRNA-S + MA10, n =7; mRNA-S + Ad5-S, n =7). **b-e**, Represented flow cytometry plot **(b,d)** and summary **(c,e)** of SARS-CoV-2 peptide-specific CD4^+^ T subsets, CD8^+^ T cell **(b,c)** at day 14 and RBD-specific B cell and plasma cell dynamic responses **(d,e)** in the BAL from different groups. **f**, SARS-CoV-2 RBD-specific IgA and IgG positive B cell dynamic responses from different groups. **g**, SARS-CoV-2 S1-specific IgA and IgG dynamic responses in the BAL from different groups. **h**, SARS-CoV-2-specific neutralizing Ab response against D614G and Omicron BA.1 in the BAL from different groups. **i**, Schematic of experimental design. *K18-hACE2* mice were immunized as indicated (Day 0, n = 3; Day 14, Omicron BA.1, n = 4; mRNA-S, n = 3; mRNA-S + Omicron BA.1, n =6; mRNA-S + Ad5-Omicron-S, n =6; Day 60, Omicron BA.1, n = 3; mRNA-S, n = 3; mRNA-S + Omicron BA.1, n =7; mRNA-S + Ad5- Omicron-S (BA.1), n =7). **j**, SARS-CoV-2 S1-specific IgA and IgG antibody dynamic responses in the BAL from different groups. **k**, Schematic of experimental design. *K18-hACE2* mice were immunized with mRNA-S, infected with BA.1 or boosted with Ad5-Omicron S (BA.1) as indicated and then challenged with SARS-CoV-2 Omicron BA2.86. **l**, Viral titers in the mouse BAL and nasal wash in different groups at 2 days post SARS-CoV-2 BA.2.86 challenge (n = 5 in each group for challenge). **c**, Two-way ANOVA with multiple-comparisons test; **l**, one-way ANOVA with multiple-comparisons test. Data presented as mean ± s.e.m. **b-h,j**, Data are representative of two independent experiments with similar results.

Consistently, upper (nasal wash) and lower airway (BAL) mucosal antigen-specific IgA levels were significantly higher in the mucosal booster group than in the hybrid group (Fig. 3g and Extended Data Fig. 5f). We observed that mucosal IgG levels, but not RBD-specific IgG^+^ B cells, were higher in boosted mice compared to those with hybrid immunity (Fig. 3 f, g). Plasma IgA, but not IgG, levels were also higher in the mucosal Ad5-S boosted group compared to those of hybrid group (Extended Data Fig. 5 g). Consequently, mucosal booster vaccination induced greatly increased respiratory nAb responses against both D614G and Omicron BA.1 viruses, which persisted for at least two months (Fig. 3h), while hybrid immunity group failed to generate detectable cross-reactive neutralizing responses against Omicron BA.1 (Fig. 3h). Nevertheless, both hybrid immunity and mucosal booster vaccination did elicit marked levels of anti-D614G and anti-Omicron nAb in the blood (Extended Data Fig. 5 h).

We also utilized K18-hACE2 transgenic mice to compare mucosal immunity induced after Omicron (BA.1) breakthrough infection with responses after Ad5-Omicron-S (BA.1) booster (Fig. 3i). In this model, we similarly observed increased cellular immune responses and increased IgA levels in the mucosal vaccination group compared to those of breakthrough infection group (Fig. 3j and Extended Data Fig. 6). To compare protective responses induced by hybrid immunity and a mucosal booster, we immunized K18-hACE2 mice with mRNA-S and infected them with Omicron BA.1 or boosted them with Ad5-Omicron-S (Fig. 3k). We then challenged the mice with Omicron BA.2.86 and examined nasal and BAL viral titers at 2 days post challenge.

As expected, mRNA-S immunization alone provided limited protection against BA.2.86 challenge (Fig. 3l), whereas hybrid immunity provided significant, albeit not sterilizing immunity against BA.2.86 (Fig. 3l). Strikingly, mucosal Ad5-Omicron S booster provided complete protection against BA.2.86 infection in both the upper and lower airways (Fig. 3l). Together, these data reveal that mucosal booster by adenoviral vaccination can generate stronger, broadly-reactive and durable respiratory mucosal immunity against SARS-CoV-2 variants compared to the hybrid immunity.

### Optimal mucosal IgA-secreting cell responses are supported by T-B interactions *in situ*

Mucosal IgA is considered to be a major determinant in protection against SARS-CoV-2 infection ^13–15^, but the underlying mechanisms regulating robust IgA production in the respiratory tract remain largely unknown. Therefore, we sought to understand the cellular and molecular mechanisms underlying enhanced IgA production by mucosal booster vaccination. We first found that RBD-specific IgA^+^ B cells and plasma cells were mainly enriched in the mucosal surface compared to those in the mediastinal lymph node (mLN), which was in sharp contrast to IgG^+^ B cells (Fig. 4a). Furthermore, respiratory IgA levels correlated with mucosal IgA-producing plasma and B cells (Fig. 4b), in contrast to mucosal IgG levels and mucosal IgG^+^ plasma cells or B cells (Fig. 4b). These data indicate that respiratory IgA is likely produced locally at the respiratory mucosa following respiratory vaccination, a notion that is also supported by other reports ^24,25^.

**Fig. 4:**
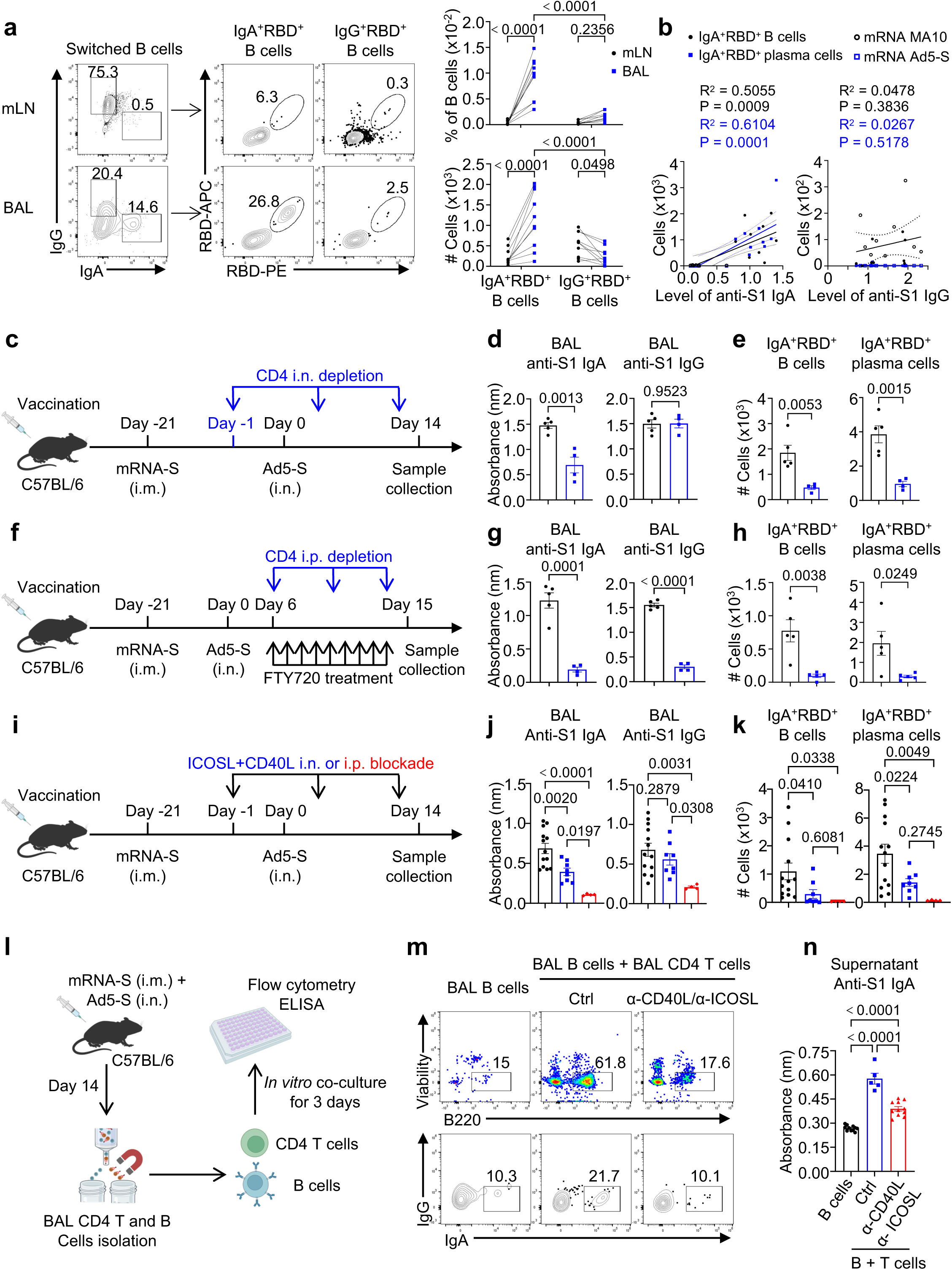
Optimal mucosal IgA responses are supported by T-B interactions *in situ*. **a,** C57BL/6 mice were immunized by one dose of mRNA-S plus Ad5-S (n = 10). Represented flow cytometry plot and summary of SARS-CoV-2 RBD-specific IgA and IgG positive B cell percentage and number in the BAL and mLN at day 14 post Ad5-S. **b,** Correlation analysis between levels of SARS-CoV-2 S1-specific IgA or IgG and SARS-CoV-2 RBD-specific IgA and IgG positive B cell and plasma cells from immunized mice as indicated at day 14 post immunization (mRNA + MA10, n = 9; mRNA + Ad5-S, n =9). **c-e**, C57BL/6 mice were immunized as indicated (Ctrl, n = 5; CD4 i.n. depletion, n =4). Schematic of experimental design **(c)**; SARS-CoV-2 S1-specific IgA and IgG responses **(d)** and SARS-CoV-2 RBD-specific IgA positive B cell and plasma cell responses **(e)** in the BAL. **f-h**, C57BL/6 mice were immunized as indicated (FTY720 treated ctrl, n = 5; FTY720 treatment plus CD4 i.p. depletion, n =4). Schematic of experimental design **(f)**; SARS-CoV-2 S1-specific IgA and IgG responses **(g)** and SARS-CoV-2 RBD-specific IgA positive B cell and plasma cell responses **(h)** in the BAL. **i-k**, C57BL/6 mice were immunized as indicated (Ctrl, n = 13; ICOSL + CD40L i.n. blockade, n =8; ICOSL + CD40L i.p. blockade, n =4). Schematic of experimental design **(i)**; SARS-CoV-2 S1- specific IgA and IgG responses **(j)** and SARS-CoV-2 RBD-specific IgA positive B cell and plasma cell responses **(k)** in the BAL. **l-n**, C57BL/6 mice were immunized as indicated (n = 5). Schematic of *in vitro* coculture experimental design **(l)**; Represented flow cytometry plot of B cell and IgA secreting cell responses **(m)** and SARS-CoV-2 S1-specific IgA responses in supernatant (B cell alone, n=11; B plus T cells, n = 5, B plus T cells with CD40L and ICOSL blockade, n = 11) **(n)**. **a**, Two-way ANOVA with multiple-comparisons test; **b**, single linear regression analysis; **d,e,g,h**, unpaired two-sided t-test; **j,k,n**, one-way ANOVA with multiple- comparisons test. Data presented as mean ± s.e.m. **a,b,j,k**, Data are pooled from two independent experiments; **d,e,g,h,n**, Data are representative of two independent experiments with similar results.

We found that respiratory IgA appeared to be CD4^+^, but not CD8^+^, T cell help dependent (Extended Data Fig. 7). The increased presence of IgA^+^ B cells (compared to IgG^+^ B cells) in the mucosal surface compared to the local draining LNs indicates that there may be local environmental factors facilitating the generation and/or expansion of IgA^+^ B cells *in situ*. To determine the roles of respiratory CD4^+^ T cells in assisting IgA production, we utilized an intranasal CD4^+^ T cell depletion strategy by intranasal (i.n.) delivery of a small amount of α-CD4, which partially depleted CD4^+^ T cells in the respiratory mucosa but not in the lymphoid tissue (Fig. 4c and Extended Data Fig. 8 a, b). Notably, respiratory CD4^+^ T cell depletion diminished BAL IgA, but not BAL IgG, nor plasma IgA and IgG production (Fig. 4d and Extended Data Fig. 8 c, d). A similar decrease in RBD-specific IgA^+^, but not IgG^+^, B cells and plasma cells was observed (Fig. 4e and Extended Data Fig. 8f) indicating that local CD4^+^ T- and B cell interaction is required for optimal IgA, but not IgG responses, as IgG is mainly produced systemically.

We complemented the strategy by blocking CD4^+^ T cell egress from secondary lymphoid organs in mice with S1P antagonist FTY720 (Fig. 4f and Extended Data Fig. 8 g-k). Ad5-S boosted mice were treated with FTY720 on day 6 post mucosal immunization, when activated T- and B cells have already migrated into the lungs from mLNs (Fig. 4f). FTY720 treatment significantly reduced T- and B cells in the blood (Extended Data Fig. 8h), indicating its effectiveness in blocking lymphocyte migration. We then depleted CD4^+^ T cells 12 hours after FTY720 injection by intraperitoneal (i.p.) delivery of α-CD4 and found that respiratory IgA levels, and IgA- expressing B and plasma cells were decreased (Fig. 4 g,h). BAL IgG levels were also decreased, likely due to CD4^+^ T cell depletion in the lymphoid organs (Fig. 4g and Extended Data Fig. 8j). Nevertheless, BAL IgG^+^ B and plasma cells were not affected by CD4^+^ T cell depletion in this scenario (Extended Data Fig. 8k).

We next systemically (i.p.) or locally (i.n.) blocked the function of CD40L and ICOS, two conventional B cell helper molecules on CD4^+^ T cells ^26^, through the injection of α-CD40L plus α-ICOS (Fig. 4i). Systemic, but not pulmonary blockade, of CD40L and ICOS function resulted in the decrease of BAL and blood S1 and RBD IgG levels (Fig. 4j and Extended Data Fig. 8l-p).

In contrast, although less effective than systemic blockade, the local inhibition of CD40L and ICOS activity also resulted in diminished IgA levels, and IgA^+^ B and plasma cells (Fig. 4j, k and Extended Data Fig. 8l-p). Altogether, these data suggest that CD4^+^ T cells provide “local” help for the optimal generation of mucosal IgA-secreting cells in the respiratory tract. To further test whether respiratory T cells could directly deliver this help to B cells in isolation, we purified BAL T- and B cells from mucosal-boosted mice and co-cultured them *in vitro* (Fig. 4l). Co-culture of respiratory T- and B cells resulted in enhanced CD40 and ICOS-dependent survival and expansion of B cells, including IgA^+^ B cells, which was accompanied by increased IgA production in the supernatant (Fig. 4l-n). These data indicate that respiratory CD4^+^ T cells exhibit “helper” function to local B cells for IgA production.

### IL-21 and Blimp-1^+^ Th1 effector cells assist mucosal IgA production

IL-21 has been shown to enhance B cell responses and antibody production ^27^. In comparison to mice immunized with mRNA alone or those with hybrid immunity, mucosal booster immunization exhibited significantly higher levels of BAL IL-21 (Fig. 5a). To assess whether mucosal IL-21 signaling is crucial for the generation of IgA-producing cells, we administered α- IL-21R intranasally to mice with mucosal booster immunization (Fig. 5b). Blocking of IL-21R in the respiratory tract resulted in a reduction of IgA^+^ B and plasma cells, along with decreased BAL IgA levels (Fig. 5c, d). Notably, blocking respiratory IL-21 signaling did not affect mucosal and systemic IgG levels (Extended Data Fig. 9). To investigate whether CD4^+^ T cell-derived IL- 21 directly promotes IgA B cell generation, we cultured mucosal T- and B cells and subsequently blocked IL-21 signaling. *In vitro* IL-21 signaling blockade diminished IgA^+^ B cell responses and IgA levels in the supernatant (Fig. 5e). These findings indicate that mucosal T cells can facilitate B cell help for IgA production, partially via IL-21 production.

**Fig. 5:**
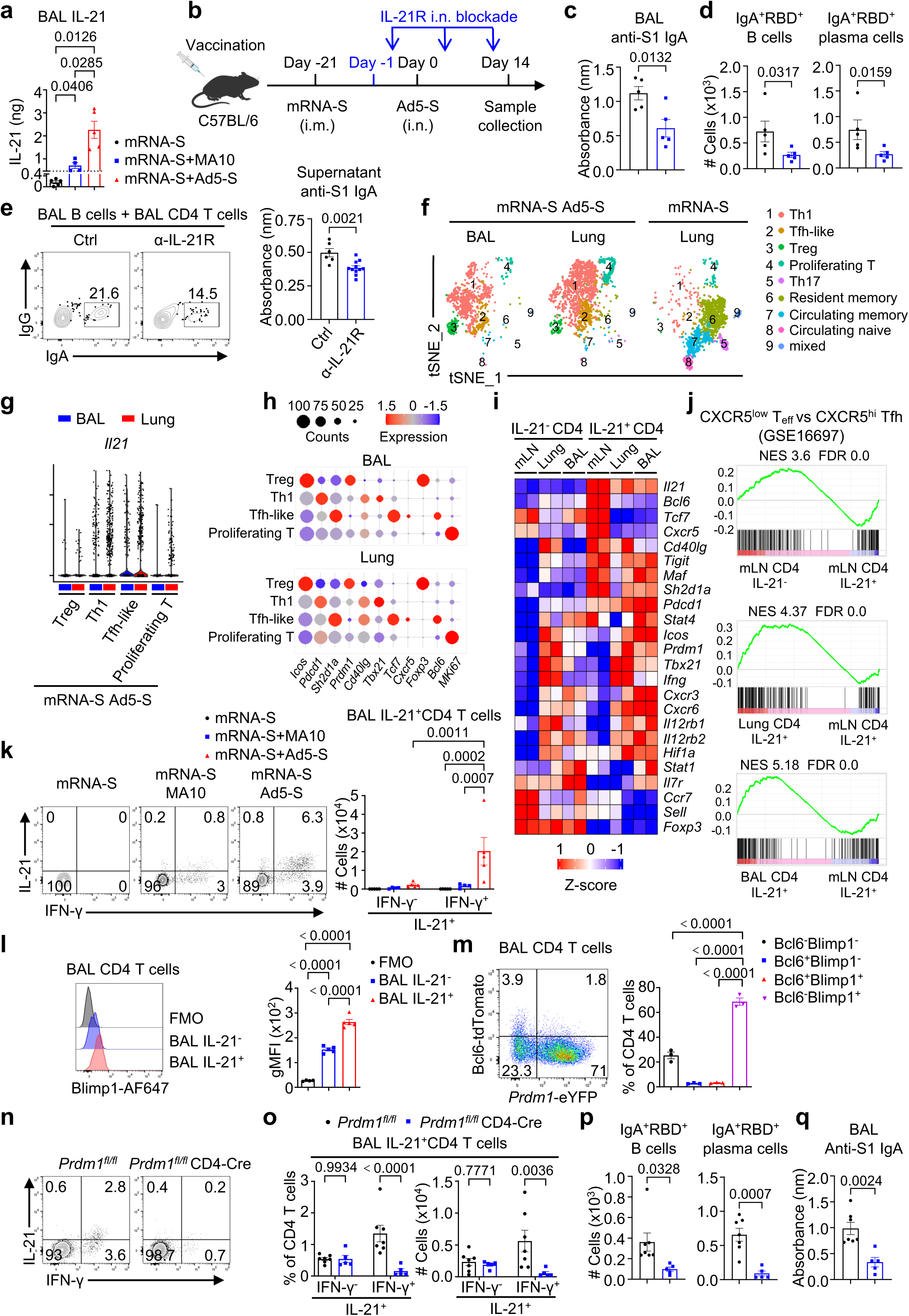
IL-21 and Blimp-1^+^ Th1 effector cells assist mucosal IgA production. **a,** Level of IL-21 in the BAL from different groups (mRNA-S, n = 5; mRNA-S + MA10, n = 4; mRNA-S + Ad5-S, n = 5) as indicated at day 6 post immunizations. **b-d**, C57BL/6 mice were immunized as indicated (Ctrl, n = 5; IL-21R i.n. blockade, n = 5). Schematic of experimental design **(b)**; SARS-CoV-2 S1-specific IgA antibody responses **(c)** and SARS-CoV-2 RBD-specific IgA positive B cell and plasma cell responses **(d)** in the BAL. **e**, C57BL/6 mice were immunized with one dose of mRNA-S plus Ad5-S (n = 5). BAL CD4^+^ T cells and B cells were isolated at day 14 for *in vitro* coculture for 3 days. Represented flow cytometry plot of IgA secreting cell response and SARS-CoV-2 S1-specific IgA responses in supernatant (B plus T cells, n = 6, B plus T cells with IL-21R blockade, n = 11). **f-h**, scRNA-seq analysis of BAL and lung parenchyma CD4^+^ T cells at day 6 post Ad5-S booster and lung parenchyma CD4 T cells without Ad5-S booster from C57BL/6 mice (mRNA + Ad5-S, n = 3; mRNA, n = 3). An UMAP plot of clustering **(f)**; *Il21* expression **(g)**, and main Th1- and Tfh-related gene expression in indicated clusters **(h)**. **i,j**, Bulk RNA-seq analysis of IL-21^+^ and IL-21^-^ CD4^+^ T cells in BAL, lung parenchyma and mLN from *IL-21-VFP* reporter mice at day 6 post Ad5-S booster (n = 5). Heatmap of Tfh- and Th1-related gene expression **(i)** and GSEA of CXCR5^low^ effector CD4^+^ T vs CXCR5^hi^ Tfh (GSE16697) in different groups compared to mLN IL-21^+^ CD4^+^ T cells **(j)**. **k**, Represented flow cytometry plot and summary of SARS-CoV-2 peptide-specific IFN-γ and IL-21 positive CD4^+^ T cells in the BAL from different groups (mRNA-S, n = 5; mRNA-S + MA10, n = 4; mRNA-S + Ad5-S, n = 5) as indicated at day 14. **l**, Represented flow cytometry plot and summary of level of Blimp-1 expression in BAL CD4^+^ T cells from mice immunized with one dose of mRNA-S plus Ad5-S at day 14. **m**, Represented flow cytometry plot and summary of percentage of Blimp-1 and Bcl6 expressions in BAL CD4^+^ T cells from Blimp-1/Bcl6 dual reporter mice immunized with mRNA-S and Ad5-S as previously at day 14 (n = 3). **n-q**, *Prdm1^fl/fl^* and *Prdm1^fl/fl^* CD4-Cre mice were immunized with mRNA-S and Ad5-S as previously (*Prdm1^fl/fl^*, n = 7; *Prdm1^fl/fl^* CD4-Cre, n = 5). Represented flow cytometry plot **(n)** and summary **(o)** of SARS-CoV-2 peptide-specific IFN-γ and IL-21 positive CD4^+^ T cells in the BAL; SARS-CoV-2 RBD-specific IgA positive B cell, plasma cell **(p)** and S1-specific IgA antibody responses **(q)** in the BAL. **a,l,m**, One-way ANOVA with multiple-comparisons test; **c-e,p,q**, unpaired two-sided t- test; **k,o**, two-way ANOVA with multiple-comparisons test. Data presented as mean ± s.e.m. **c- e,k,l**, Data are representative of two independent experiments with similar results.

To understand the molecular signature of those mucosal helper CD4^+^ T cells, we performed scRNA-seq on purified lung CD4^+^ T cells from mRNA-S immunized mice, as well as lung and BAL CD4^+^ T cells from mRNA-S immunized mice that were additionally boosted with Ad5-S. We did not perform scRNA-seq on BAL T cells from mice immunized with mRNA alone, as these mice had very few CD4^+^ T cells in the BAL. Clustering analysis revealed that BAL and lung CD4^+^ T cells from mucosal-boosted mice predominantly consisted of Th1 effector cells, which express *Tbx21* (T-bet) and *Prdm1* (Blimp-1) (Fig. 5f and Extended Data Fig. 10), with smaller populations of regulatory T (Treg) (expressing *Foxp3*), Tfh-like (expressing *Bcl6*), proliferating (expressing *Mki67*), and other cells. Cytokines are typically challenging to detect by scRNAseq. Notably, in addition to the minor Tfh-like cell population, we detected IL-21 in BAL and lung Th1 cells (Fig. 5g and Extended Data Fig. 10d). Furthermore, these BAL and lung Th1 cells expressed key B cell helping molecules, including *Cd40lg* and *Icos,* which are important for mucosal IgA production (Fig. 4i-n), as well as *Pdcd1, Sh2d1a* among others (Fig. 5h).

Given the limited sequencing depth of scRNA-seq, we conducted total RNA-seq on sorted IL- 21-VFP^+^ or IL-21-VFP^-^ CD4^+^ T cells isolated from the mLN, lung, and BAL of mRNA plus Ad5-S immunized IL-21-VFP reporter mice (Extended Data Fig. 11a-c). DEG and Gene Set Enrichment Analysis (GSEA) revealed that BAL and lung IL-21^+^ cells were enriched with genes associated with CD4^+^ effector T cell differentiation, interferon response, KRAS signaling and STAT3 signaling etc. compared to mLN IL-21^+^ cells, which were Tfh cells (Fig. 5i, j and Extended Data Fig. 11d, e). Additionally, IL-21^+^ CD4^+^ T cells in BAL and lung showed reduced expression of lymphoid Tfh cell-associated genes such as *Bcl6, Tcf7,* and *Cxcr5*, but increased levels of terminal Th1-associated genes including *Prdm1* (Blimp-1), *Tbx21, Ifng,* and *Cxcr3* (Fig. 5i and Extended Data Fig. 11d). These cells also expressed comparable or slightly lower levels of critical B cell helping molecules such as *CD40lg, Icos, Il21, Sh2d1a*, etc., compared to IL-21 producing cells in the mLN (Fig. 5i). Importantly, these IL-21^+^ cells did not express *Foxp3*, unlike IL-21^-^ cells, indicating their effector lineage (Fig. 5i).

The fact that mucosal IL-21-producing cells co-expressed high levels of B cell helper molecules and were enriched with effector Th1 cell gene signatures, including Blimp-1 expression, was unexpected, as Blimp-1 is widely believed to repress B cell helper function ^28,29^. To this end, flow cytometry analysis confirmed that most IL-21-producing CD4^+^ T cells co-expressed IFN-γ, affirming their Th1 lineage (Fig. 5k). Notably, IL-21-producing cells expressed higher levels of Blimp-1 compared to IL-21 negative cells (Fig. 5l). Blimp-1 has been shown to program peripheral homing terminal effector T cells. Consistent with this, using Blimp-1/Bcl6 double reporter mice, we verified that most BAL T cells are Blimp-1^+^ rather than Bcl6^+^, a marker of B cell helping Tfh or Tfh-like cells (Fig. 5m) ^28–31^.

To directly address whether Blimp-1^+^ Th1 cells assist mucosal IgA responses, we generated bone marrow (BM) chimeric mice composed of *Prdm1^fl/fl^* or T cell-specific Blimp-1-deficient (*Prdm1^fl/fl^* CD4-Cre) BM cells. We then immunized the mice with mRNA-S plus Ad5-S. Consistent with the roles of Blimp-1 in Th1 effector differentiation and peripheral homing, Blimp- 1 deficiency reduced respiratory IFN-γ-producing CD4^+^ T cells (Fig. 5n). Additionally, Blimp-1 deficiency eliminated IL-21^+^ IFN-γ^+^ cells, but not IL-21^+^ IFN-γ^-^ cells, supporting the idea that Blimp-1 is necessary for the mucosal generation or homing of these IL-21-producing Th1 cells (Fig. 5o). Consequently, mucosal IgA^+^ B and plasma cells were significantly diminished following CD4^+^ T cell Blimp-1 deficiency (Fig. 5p). Moreover, mucosal IgA levels, and to a lesser extent, IgG levels, were decreased after Blimp-1 ablation in T cells (Fig. 5q and Extended Data Fig. 12). Collectively, these data indicate that respiratory Blimp-1^+^ IL-21-producing effector Th1 cells support local IgA-producing B cell accumulation and IgA responses in the respiratory tract.

### TGF-β and lung macrophages facilitate robust mucosal IgA responses after mucosal immunization

TGF-β is a critical cytokine for promoting IgA class switching in the gut ^32^. We found that mucosal adenoviral immunization led to an increase BAL TGF-β levels, higher than mRNA vaccination alone or hybrid immunity (Fig. 6a). Importantly, BAL B cells expressed higher levels of TGFBRII than those of mLN B cells (Fig. 6b). To examine the roles of TGF-β signaling in promoting the development of IgA^+^ B cells, we generated mixed BM chimeric mice by injecting irradiated mice with a mixture of BM from CD45.1^+^ control and CD45.1^+^ CD45.2^+^ TGFBRII inducible knockout mice (*Tgfbr2*-iKO). We then immunized the mice with mRNA, injected tamoxifen to delete TGFBRII in *Tgfbr2*-iKO cells, and followed with Ad5-S immunization (Fig. 6c and Extended Data Fig. 13 a, b). TGFBRII deficiency resulted in greatly diminished IgA^+^ B and plasma cells, suggesting B cell-intrinsic TGF-β signaling is required for IgA-secreting cell generation (Fig. 6d, e). To examine the specific function of respiratory TGF-β in IgA responses, we blocked local TGF-β in the mucosal immunization model through i.n. delivery of α-TGF-β (Fig. 6f). Strikingly, pulmonary TGF-β blockade resulted in decreased RBD-specific IgA^+^ B and plasma cells, which were accompanied by diminished antigen-specific IgA levels (Fig. 6 g, h), but not antigen-specific IgG or IgG-producing B cells (Extended Data Fig. 13 c-f).

**Fig. 6:**
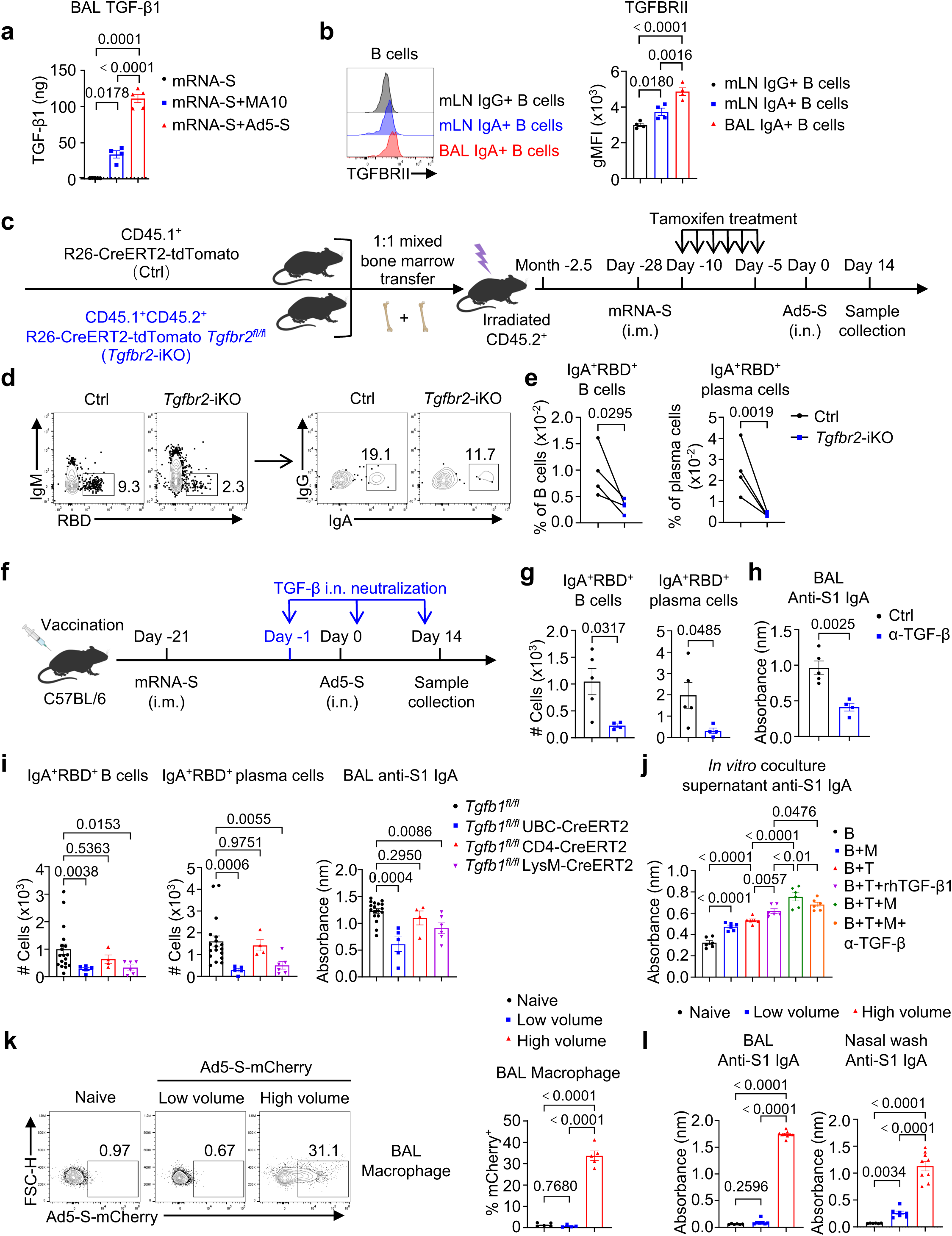
Lung macrophage and TGF-β assist optimal mucosal IgA production. **a,** Level of TGF-β1 in the BAL from different groups (mRNA-S, n = 5; mRNA-S + MA10, n = 4; mRNA-S + Ad5-S, n = 5) as indicated at day 6 post immunizations. **b**, Represented flow cytometry plot and summary of level of TGFBRII expression in IgA^+^ B cells in the BAL and IgA^+^ B and IgG^+^ B cells in mLN from mice immunized with mRNA-S and Ad5-S as previously at day 14 (n = 4). **c-e,** Mixed CD45.1^+^R26-CreERT2-tdTomato (Ctrl) and CD45.1^+^CD45.2^+^R26- CreERT2-tdTomato *Tgfbr2^fl/fl^* (Tgfbr2-iKO) BM (1:1) chimeric mice were immunized and treated as indicated (n = 4). Schematic of experimental design **(c)**; Represented flow cytometry plot **(d)** and summary **(e)** of SARS-CoV-2 RBD-specific IgA positive B cell and plasma cell percentage. **f-h**, C57BL/6 mice were immunized as indicated (Ctrl, n = 5; TGF-β i.n. blockade, n = 4). Schematic of experimental design **(f)**; SARS-CoV-2 RBD-specific IgA positive B cell, plasma cell **(g)** and S1-specific IgA antibody responses **(h)** in the BAL. **i**, SARS-CoV-2 RBD-specific IgA positive B cell, plasma cell and S1-specific IgA responses in the BAL from indicated mice immunized with mRNA-S and Ad5-S as previously at day 14 (*Tgfb1^fl/fl^*, n = 11; *Tgfb1^fl/fl^* UBC- CreERT2, n = 5; *Tgfb1^fl/fl^* CD4-CreERT2, n = 4; *Tgfb1^fl/fl^* LysM-CreERT2, n = 6). **j**, C57BL/6 mice were immunized with one dose of mRNA-S plus Ad5-S (n = 5). BAL CD4^+^ T, B cells and macrophages were isolated at day 14 for *in vitro* coculture for 3 days. SARS-CoV-2 S1-specific IgA responses in supernatant (n = 5 for each group). **k**, C57BL/6 mice were immunized with Ad5-S-mCherry (Naïve, n = 5; low volume, n = 4; high volume, n = 5). Represented flow cytometry plot and summary of mCherry^+^ macrophage percentage in the BAL. **l**, SARS-CoV-2 S1-specific IgA responses in the BAL and nasal wash from mice immunized with mRNA-S and Ad5-S as previously at day 14 (Naïve, n = 6; low volume, n = 7; high volume, n = 9). **a,b, i-l**, One-way ANOVA with multiple-comparisons test; **e**, paired two-sided t-test; **g,h**, unpaired two- sided t-test. Data presented as mean ± s.e.m. **b,d-h,j,k**, Data are representative of two independent experiments with similar results. **i**, Data are pooled from three independent experiments.

TGF-β is ubiquitously expressed by many cells ^33^.T cell-derived TGF-β in particular, has been shown to promote gut IgA production ^32,34^. To understand the cellular source of TGF-β required for respiratory IgA production, we crossed Tgfb1^fl/fl^ mice with UBC-CreERT2, CD4-CreERT2 or LysM-CreERT2 mice to generate inducible TGF-β-deficient mice in all cells, CD4^+^ T cells and myeloid cells respectively. We immunized these mice with mRNA plus Ad5-S and found that the depletion of TGF-β in myeloid cells, but not in T cells, during the mucosal immunization phase caused reduced levels of IgA^+^ B and plasma cells, as well as diminished antigen-specific IgA (Fig. 6i). Similarly, Treg-specific deficiency of TGF-β also did not affect BAL IgA levels (Extended Data Fig. 14 a-c). Notably, among myeloid cells, resident alveolar macrophages (AMs) are a major source of TGF-β1 ^35^. Indeed, we found that the depletion of AMs following diphtheria toxin (DTX) treatment in CD169-DTR mice resulted in diminished IgA^+^ B and plasma cells, indicating that AMs are required for optimal respiratory IgA-producing B cell generation (Extended Data Fig. 14 d, e). To directly examine whether AMs can help B cell IgA production through TGF-β1, we utilized an *in vitro* co-culture system by adding AMs to the BAL CD4^+^ T cell and B cell culture (Extended Data Fig. 14f). Inclusion of AMs in the culture promoted IgA^+^ B cell generation and IgA secretion, which was at least partially dependent on TGF-β1 (Fig. 6j and Extended Data Fig. 14g). These data reveal that lung macrophages, likely AMs, promote mucosal IgA production through TGF-β. Together, our results suggest that a local cellular network involving lung macrophages, periphery-homing terminal Th1 cells, and B cells cooperate to promote optimal respiratory mucosal IgA production following mucosal immunization.

### Lower airway adenovirus delivery is necessary for robust upper and lower airway IgA

The data above strongly suggest that an optimal immunization strategy for inducing robust respiratory mucosal IgA requires the involvement of AMs, which reside in the lung (lower airway). Therefore, we hypothesized that lower airway delivery of replication-deficient adenovirus, which are widely used in immunization strategies due to safety considerations of using replication-competent virus, would be necessary to induce respiratory mucosal IgA responses since the vector would not reach the distal lungs without replication, upon delivery to the upper respiratory tract. To test this hypothesis, we intranasally delivered a low volume (10 µl, so that the virus mainly stay in the nasal cavity and the upper airway) or our regular delivery volume (high volume, 30 µl, so that the virus can reach to the lower airway and the lung from the nasal cavity) of the same viral titer of Ad5-S-mCherry virus into the mice. We confirmed that both low and high volumes of adenovirus administration induced innate inflammatory responses in the nasal cavity (Extended Data Fig. 15 a, b). However, only the high volume of adenovirus administration was able to deliver the virus to AMs, as indicated by mCherry expression (Fig. 6k), reflecting that high volume intranasal administration resulted in efficient delivery to the lower airways. Strikingly, high volume delivery not only generated robust mucosal IgA in the lower airway (BAL), but also resulted in much higher levels of IgA and IgG in the upper respiratory tract (nasal wash) (Fig. 6m and Extended Data Fig.15 c-e). These data together suggest that the delivery of adenoviral constructs to the lower airways is crucial for effective mucosal immunization thus the induction of strong mucosal immunity throughout the respiratory tract.

## Discussion

The limited efficacy of mRNA vaccines in inducing respiratory humoral and cellular immunity highlights the need for developing mucosal SARS-CoV-2 booster vaccines. Notably, adenovirus-based mucosal booster vaccine candidates have demonstrated robust respiratory mucosal immunity in pre-clinical studies, eliciting significant IgA and CD8^+^ T cell responses in various animal models, including non-human primates ^36–39^ ^40^. These responses have been shown to confer superior protection against SARS-CoV-2 variant challenges compared to systemic immunization ^36–40^. Real-world epidemiological data also suggest that adenovirus-based mucosal booster vaccines elicit strong anti-SARS-CoV-2 immune responses and offer greater protection against breakthrough infections from variants compared to inactivated virus vaccines, lasting more than 10 months ^38,41–45^. These clinical and preclinical findings underscore the potential of mucosal vaccine boosters, including adenovirus-based strategies. However, unlike the results from inhaled or intratracheal delivery of the adenovirus vaccine, a nasal-spray adenovirus vaccine candidate was ineffective in inducing strong respiratory humoral responses in humans ^46^. While the reasons for these discrepancies are still not fully understood, our results offer a plausible explanation: a nasal spray vaccine may not stimulate full lung macrophage responses, which are necessary for inducing robust IgA-mediated mucosal immunity. We speculate that strategies capable of targeting lung macrophages and inducing stronger “helper” responses via respiratory tract-homing Th1 cells could be key to the efficacy of future mucosal vaccines.

Interactions between T and B cells in lymphoid tissues are crucial for developing humoral and cellular immunity against viruses. Class-switched high-affinity Ab production requires CD4^+^ T cells, particularly follicular helper T cells (Tfh) in the germinal centers (GCs). The transcription factor Bcl6 is a master regulator for the development of Tfh or peripheral Tfh-like cells^28,30,47^ and is necessary for T cell help to B cells in secondary lymphoid organs ^26,28,30,47–51^. Recent reports have suggested that Tfh and GC reactions are required for the generation of T cell-dependent IgA-secreting cells in the gut and the upper respiratory tract ^16,52^. In contrast, Blimp-1, which opposes Bcl6 expression and function, is generally believed to promote terminal effector T cell differentiation and inhibit B cell help from T cells ^28,47,53,54^. However, these conclusions were drawn mostly from studies conducted in secondary lymphoid organs during primary infection or immunization, focusing on systemic Ab responses. Our results indicate that following initial activation and programming of IgA^+^ B cells in the regional lymphoid organs, periphery-homing Blimp-1^+^ IL-21-expressing Th1 cells also play an important role in facilitating robust respiratory mucosal IgA^+^ B and plasma cell responses in the context of a respiratory booster vaccination. Nevertheless, it remains unclear whether Blimp-1 directly promotes IL-21 and other B cell helper molecules in respiratory Th1 cells or functions by modulating the migration, differentiation, and/or expansion of these cells.

Reports have suggested that IL-21 and particularly TGF-β could directly induce B cell class switch recombination (CSR) and promote IgA production ^55^. It is therefore tempting to speculate that Th1 cell-derived IL-21 and macrophage-derived TGF-β may directly promote B cell class switching at the mucosal surface *in situ.* Future studies are warranted to dissect the exact roles of these Th1 effector cells and macrophages in assisting IgA-secreting cells with their migration, CSR, proliferation, and/or survival *in vivo.* Nevertheless, our data and recent reports support a “two-step” model regarding the generation of full magnitude of respiratory mucosal IgA- secreting cells during respiratory vaccination (Extended Data Fig.16). In this model, the initial activation and class-switching of IgA-secreting cells are programmed at regional lymphoid tissues, followed by further assistance from mucosa-residing effector T cells and local macrophages to facilitate the full magnitude of IgA-producing cell responses.

Unlike intramuscular vaccination, SARS-CoV-2 infection induces detectable respiratory mucosal humoral and cellular immunity ^3^. Natural immunity developed after infection has been widely considered an effective way to protect against infection and/or disease development during secondary infection ^7,8^. In the context of SARS-CoV-2, vaccination plus infection can further boost the quantity and quality of humoral and cellular immunity in the circulation and the oral mucosa ^9–12^. Our data and others further show that hybrid immunity is characterized by increased respiratory humoral responses compared to vaccination or infection alone ^56^.

However, whether a mucosal vaccine can elicit stronger, broader, and durable immune responses to confer superior mucosal protection against SARS-CoV-2 compared to hybrid immunity has become a key question for justifying the development of future mucosal vaccines. Our results that the mucosal booster can elicit higher cross-reactive and durable mucosal Ab levels, including IgA, as well as more robust local T cell responses compared to natural infection or hybrid immunity, provide important basis for the development of optimal mucosal vaccines against SARS-CoV-2 (Extended Data Fig.16). The relatively low levels of mucosal responses in hybrid immunity, compared to those elicited by a booster adenoviral vaccination, can be partially attributed to suboptimal IL-21-producing Th1 responses and insufficient local TGF-β production. This may be due to the limited spread of SARS-CoV-2 infection to the lungs, and/or a shortened infection duration in vaccinated individuals, particularly in the case of the Omicron lineage known to preferentially infect the upper airway ^21^.

In summary, our study renews calls for the development of mucosal vaccines in the future that can induce optimal respiratory protective immunity to impede SARS-CoV-2 infection at the primary viral entry site, hence preventing the subsequent development of acute and/or chronic disease (long COVID) ^57,58^. Our characterization of the coordinated interactions between lung macrophages, effector T cells, and B cells, that promote robust upper and lower mucosal IgA responses aids in innovative vaccine strategies and therapeutic approaches to elicit strong mucosal immunity against SARS-CoV-2 and other respiratory pathogens.

## Methods

### Study cohorts

The study sample included children with difficult asthma living in a mixed urban/rural region in central Virginia to a university-based program. Criteria to confirm asthma included recurrent or daily cough and wheeze, physician diagnosis, treatment with any asthma medication, and/or airflow limitation (FEV_1_/FVC < 0.85) with spirometry. Children were treated according to standard guidelines ^59^ and reevaluated every six months. Treatment was adjusted according to exacerbation history, lung function results, and symptom scores. Adherence to treatment was monitored according to pharmacy refill records ^60^. Criteria for bronchoscopy included treatment failure despite adequate adherence with any one of the following: two or more episodes of cough and wheeze per year which required oral prednisone, two or more urgent care visits for wheeze, persistently elevated symptoms based on an asthma control test value < 20 units, or persistent airflow limitation. Caregivers provided history of previous SARS-CoV-2 infections, dates, and dates of the last mRNA vaccination for SARS-CoV-2 as confirmed by the medical record. The legal guardians and caregivers of children provided informed consent and older children assent as approved by the institutional review board (IRB-HSR # 18975). The study sample was characterized at enrollment according to SARS-CoV-2 history and vaccine status into four categories: uninfected/unvaccinated (control), uninfected vaccinated (vaccine), infected/unvaccinated (COVID), and infected/vaccinated (hybrid). However, measures of the N- IgG antibody OD450 in BAL (1:5) and plasma (1:25) showed that children with no history of past SARS-CoV-2 infection or vaccination had elevated BAL and plasma N-IgG titers consistent with past infection. Accordingly, children with BAL + Plasma IgG OD values > 0.5 were re-assigned to accurate categories. Detailed information was shown in Extended Data tables 1 and 2.

### Human sample collection

Characterization procedures have been published in detail ^61,62^. Data were stored in a secured bioregistry that included demographics, anthropometrics, symptom history, comorbidities, treatment details, survey of social determinants, and environmental exposures. The evaluation included detailed examination of the upper and lower respiratory tract, lung lavage (BAL) nucleic acid multiplex for respiratory viruses, bacterial cultures, granulocyte differential count, and blood for type 2 inflammatory markers, IgE directed against specific allergens, and high-sensitivity-C-reactive protein (CRP). BAL granulocyte categories were defined according to the studies of Hastie et al ^62^. Lung function was measured in children over four years of age with standard methods and interpreted according to international reference standards.

Bronchoscopy was accomplished with general anesthesia (sevoflurane/propofol) administered by a pediatric anesthesiologist via a laryngeal mask airway. BAL specimens were obtained from the right middle lobe through the instillation one cc per kg (max 30 cc) sterile 0.9% saline warmed to 37°C, limited to three flushes, with an average return of 0.28 fluid per instill ate. BAL was processed immediately unfiltered, spun at 600g for 5 minutes with the supernatant frozen at -80°C for analysis. Nasal wash is collected from 1.5 ml phosphate buffered saline instilled via an intranasal atomizer (MAD, Cardinal Heath) into each nasal cavity, then aspirated (BBG, Philips Medical Systems) into a trap containing 2 ml RPMI nutrient ^63^, followed by processed at 600 g for 5 minutes with the supernatant frozen at -80°C for analysis. A sample of venous blood was collected, spun, and the plasma was frozen for analysis.

### Mice

WT CD45.2 C57BL/6 (#000664), CD45.1^+^ C57BL/6 (#002014), K18-hACE2 (#034860), IL-21- VFP reporter (#030295), *Prdm1*-eYFP (#008828), Rosa26-eYFP (007903), *Prdm1^fl/fl^* (#008100), *Tgfb1^fl/fl^* (#065809), CD4-Cre (#022071), Foxp3-YFP-Cre (#016959), CD4-CreERT2 (#022356), UBC-CreERT2 (#007001), LysM-CreERT2 (#032291), and Rosa26-CreERT2-tdTomato (#008463) mice were originally purchased from the Jackson Laboratory. CD169-DTR mice were originally from Dr. Masato Tanaka ^64^. *Tgfbr2^fl/fl^* mice were originally from Dr. Stefan Karlsson ^65^. Bcl6-tdTomato reporter mice were originally from Dr. Yohsuke Harada ^66^, *Tgfb1^exon2fl/fl^*mice were generated at The Ohio State University Comprehensive Cancer Center ^67^. *Prdm1^fl/fl^* mice were bred with CD4-Cre to generate *Prdm1^fl/fl^* CD4-Cre mice. Rosa26-CreERT2-tdTomato mice were bred with CD45.1^+^ C57BL/6 congenic mice to generate CD45.1^+^ Rosa26-CreERT2-tdTomato.

*Tgfbr2^fl/fl^* mice were bred with Rosa26-CreERT2-tdTomato and CD45.1^+^ C57BL/6 congenic mice to generate CD45.1^+^CD45.2^+^ *Tgfbr2^fl/fl^* Rosa26-CreERT2-tdTomato. *Tgfb1^fl/fl^* mice were bred with UBC-CreERT2 or LysM-CreERT2 mice to generate *Tgfb1^fl/fl^* UBC-CreERT2 or *Tgfb1^fl/fl^* LysM- CreERT2 mice. *Tgfb1^fl/fl^* mice were bred with CD4-CreERT2 and Rosa26-eYFP mice to generate *Tgfb1^fl/fl^* CD4-CreERT2 mice. *Tgfb1^exon2fl/fl^* mice were bred with Foxp3-YFP-Cre mice to generate *Tgfb1^fl/fl^* Foxp3-Cre mice. Experiments were conducted using age and sex matched male and female mice at 8-14 weeks of age. All animals were maintained in specific pathogen– free barrier facilities, used in accordance with protocols approved by the Institutional Animal Care and Use Committees of the University of Virginia (Charlottesville, VA, #4369 and #4370) and followed all relevant ethical regulations.

### Vaccination and infection

mRNA-S for most experiments were made as previously reported ^3^. Briefly, antigens encoded by the mRNA vaccines were derived from SARS-CoV-2 isolate Wuhan-Hu-1 (GenBank MN908947.3). Nucleoside-modified mRNAs encoding the full length of the Spike protein from SARS-CoV-2 with two proline mutations (mRNA-S) were synthesized by *in vitro* transcription using T7 RNA polymerase (MegaScript, Ambion). mRNAs were formulated into lipid nanoparticles (LNPs) using an ethanolic lipid mixture of ionizable cationic lipid and an aqueous buffer system. Formulated mRNA-LNPs were prepared according to RNA concentrations (∼1 μg/μl) and were stored at −80°C for animal immunizations. 1 μg of mRNA-S in 50 μl PBS was used to intramuscularly immunize mice with one or two doses in most experiments. Bivalent mRNA-S from Moderna was purchased from Pharmacy at the Virginia Commonwealth University (Richmond, VA). 2 μg of bivalent mRNA-S in 50 μl PBS was used to intramuscularly immunize CD196-DTR, DTR-littermate control mice and C57BL/6 mice for *in vitro* experiments with one dose. 5 × 10^8^ plaque-forming units (PFU) of Ad5-S, Ad5-Omicron-S and Ad5-S- mCherry (University of Iowa Viral Vector Core) normally in 30 μl naïve Dulbecco’s Modified Eagle Medium (DMEM) (10 μl for small volume) were used to intranasally immunize mice after being anesthetized by intraperitoneal injection of ketamine and xylazine. 1.2 × 10^5^ PFU SARS- CoV-2 MA10 strain (received from Dr. Barbara J Mann), 3.5 × 10^4^ PFU Omicron BA.1 strain (received from Dr. Barbara J Mann) in 50 μl DMEM were used to intranasally immunize mice after being anesthetized by intraperitoneal injection of ketamine and xylazine. 2 × 10^4^ PFU Omicron BA.2.86 strain (from BEI resources) in 75 μl DMEM were used to intranasally challenge mice after being anesthetized by intraperitoneal injection of ketamine and xylazine. Infected mice were monitored daily for weight loss and clinical signs of disease for 2 weeks, following twice a week for the duration of the experiments. The mortality rate of mice calculated as “dead” were either found dead in cage or were euthanized as mice reached 70% of their starting body weight which is the defined humane endpoint in accordance with the respective institutional animal protocols. At the designated endpoint, mice were humanely euthanized by ketamine/xylazine overdose and subsequent cervical dislocation. All work with Ad5 immunization was approved under Animal Biosafety Level 2/Biosafety Level 2 conditions and was performed with approved standard operating procedures and safety conditions by the UVA Institutional Biosafety Committee. All work with SARS-CoV-2 infection was approved under Animal Biosafety Level 3/Biosafety Level 3 conditions and was performed with approved standard operating procedures and safety conditions by the UVA Institutional Biosafety Committee.

### Mouse sample collection

Mice were injected intravenously with 1.5 μg of α-CD45 (1.5 μg/ mouse in 100 μl PBS) labeled with various fluorochromes. Two minutes after injection, mice were euthanized with an overdose of ketamine/xylazine and processed 3 min later. Nasal wash, BAL, blood, Lung and mLN were collected for analysis. Isolated plasma inactivated for 30 min at 56°C, supernatant of 1.6 ml BAL fluid, and 500 μl of nasal wash were collected and stored at −80°C for ELISA or neutralization assay. BAL and mLN single cell suspensions were passed once through 70-μm cell strainers (Falcon) before proceeding to the next step. To prepare single cells from lung tissue, lung was cut into small pieces, digested with type 2 collagenase (Worthington Biochemical), and dissociated in 37°C for 30 min with Gentle-MACS (Miltenyi Biotec). Cells were further ground through 70-μm cell strainer and washed with plain Iscove’s Modified Dulbecco’s Medium (IMDM) (Gibco). After red blood cell lysis, cells were centrifuged and resuspended in completed IMDM (10% fetal bovine serum (FBS), along with 1% of penicillin-streptomycin and L-glutamine) for flow cytometry analysis.

### *In vivo* drug treatment

For respiratory CD4^+^ T cell depletion, mice were euthanized with ketamine/xylazine and intranasally injected with 10 μg of α-CD4 (GK1.5, BioXcell) in 75 μl every six days starting one day before Ad5-S immunization. For CD4^+^ T cell depletion intranasally, mice were injected with 10 μg of α-CD4 (GK1.5, BioXCell) in 75 μl every six days starting at one day before Ad5-S immunization. For systemic CD4^+^ T cell depletion, mice were intraperitoneally injected with 250 μg of α-CD4 every six days starting one day before or six days after Ad5-S immunization.

ICOSL/CD40L systemic blockade was achieved by the intraperitoneally injection of 250 of α-ICOSL (HK5.3, BioXCell) and α-CD40L (MR-1, BioXCell) every six days starting one day before Ad5-S immunization. For respiratory blockade, mice were intranasally injected with 100 μg of α-ICOSL and α-CD40L in 50 μl for the first dose followed with 50 μg of α-ICOSL and α-CD40L in 50 μl every six days starting one day before Ad5-S immunization. For respiratory IL-21R blockade or TGF-β neutralization, mice were intranasally injected with 100 μg of α-IL-21R (4A9, BioXCell) or α-TGF-β (1D11.16.8, BioXCell) in 50 μl for the first dose followed with 50 μg of α- IL-21R or α-TGF-β in 50 μl every six days starting one day before Ad5-S immunization. Isotype control immunoglobulin G (IgG) was used as controls. In some experiments, FTY720 (1 mg/kg; Cayman) was administrated via intraperitoneal injection daily from day 6 post Ad5-S immunization to block lymphocyte migration until mouse euthanasia. To induce gene recombination in mice with CreERT2, tamoxifen (2 mg/mice/day, Sigma-Aldrich) was diluted in warm sunflower oil (Sigma-Aldrich) and daily treated via intraperitoneal route for six consecutive times. CD169-DTR mice and DTR-littermate controls were intraperitoneally injected with diphtheria toxin (300 ng/mouse, Sigma,) every 3 days from the time indicated.

### Virus titer measurement

Viral titers were determined using plaque assay in the BAL and nasal wash collected from mice at day 2 post challenge. Vero E6 cells were seeded in 12-well plates 1 day prior to infection, and then 10-fold serial dilutions of samples were added on cell monolayers. The plates were incubated at 37°C and 5% CO_2_ for 2 hour, shaking every 15 minutes. After adsorption, cells were overlaid with 2 ml Avicel overlay, a 1:1 mixture of 2.4% Avicel and 2 x DMEM supplemented with 2% FBS. Plates were fixed on day 5 with 4% paraformaldehyde and stained with 0.1% crystal violet solution. Plaques were manually counted and PFUs were calculated according to the following equation: average # of plaques/dilution factor × volume of diluted virus added to the well.

### Cell isolation and *in vitro* co-culture

To isolate CD4^+^ T, B cells and macrophages, single cell suspension of the BAL was generated from mRNA-S plus Ad5-S as described above and isolated with CD4-, B220- or SiglecF- microbeads (Miltenyi Biotec) according to the manufacturer’s instructions. Purified cells were seeded in completed DMEM in 96-well (30,000 per cell type) plates and incubated for 3 days at 37°C 5% CO_2_ with Spike_62-76_ peptide (2 μg/mL) and α-IgM (20 μg/mL). Supernatant and cell pellets were collected for IgA antibody measurement and IgA-secreting cell analyses, respectively. In some experiments, α-ICOSL/ α-CD40L (20 μg/ml), α-IL-21R (20 μg/ml), α-TGF-β (50 μg/ml), or TGF-β (50 ng/ml) were added in the system.

### Flow cytometry analysis

As previously reported ^3^, cells from tissue were stained with antibodies as listed in Extended Data table 3. All the surface staining was done on ice for 30 mins except for CXCR5 staining, for which α-CXCR5-Biotin was stained for 1 h, followed by staining with Brilliant Violet 421-labeled streptavidin on ice for 30 mins. Intracellular cytokine staining was performed to detect SARS-CoV-2-specific T cell response. Briefly, cells were washed with FACS buffer and resuspended with complete DMEM with 10 mM Hepes supplemented with 10% FBS, 2-mercaptoethanol, sodium pyruvate, nonessential amino acids, penicillin-streptomycin, and l-glutamine. Cells were then stimulated with 2 μg/mL Spike_62-76_ peptide (GenScript) and 1 μg/mL Spike_539-546_ peptide (GenScript) for stimulation for 6 hours. In the last 2 hours of incubation, protein transport inhibitor brefeldin A was added. After stimulation, cells were first stained for surface markers on ice for 30 mins. After washing with PBS, cells were resuspended with Zombie dye for viability staining and incubated at room temperature (RT) for 10 mins. After surface and viability staining, cells were fixed with fixation buffer (BioLegend) and permeabilized with Perm/Wash buffer (BioLegend), followed by intracellular cytokine staining on ice for 30 mins. Cells were then washed twice with Perm/Wash buffer and resuspended with FACS buffer. For IL-21 staining, as previously ^68^, IL-21R-Fc chimera protein was added for 1 h at RT, followed by staining with AF647-labeled α-Human IgG on ice for 30 mins. To detect RBD-specific B cells, we incubated recombinant RBD proteins coupled with phycoerythrin (PE) and allophycocyamin (APC) with the cells for 40 mins at RT. RBD-PE and RBD-APC double-positive B cells were identified as RBD^+^ B cells. To detect S_539–546_ epitope–specific CD8 T cells, we incubated SARS-CoV-2 S_539–546_ major histocompatibility complex class I tetramer (H-2Kb) (National Institutes of Health (NIH) Tetramer Core) with the cells for 30 mins at 4°C. To stain Blimp-1, after surface and viability staining, cells were fixed with fixation buffer (Cytek Biosciences) and permeabilized with Perm buffer (Cytek Biosciences), followed by staining at RT for 1 h. After staining, cells were washed twice with Perm/Wash buffer (BioLegend) to reduce the background. Cell population data were acquired on Attune NxT (Thermo Fisher Scientific) and analyzed using FlowJo Software (10.8.1, Tree Star Inc.).

### Construction of bone marrow chimeras

For Bcl6-tdTomato/*Prdm1*-eYFP, *Prdm1^fl/fl^* CD4-Cre, *Prdm1^fl/fl^*, *Tgfb1^fl/fl^* Foxp3-Cre and WT ctrl for *Tgfb1^fl/fl^* Foxp3-Cre mice, Bone marrow cells from each were transferred to the irradiated (1,100 rad) WT recipient mice to create chimeric mice. Additionally, Bone marrow cells from CD45.1^+^ Rosa26-CreERT2-tdTomato and CD45.1^+^CD45.2^+^ *Tgfbr2^fl/fl^* Rosa26-CreERT2- tdTomato mice were mixed at a ratio of 1:1, the mixed bone marrow were then transferred to the irradiated (1,100 rad) WT CD45.2^+^ recipient mice to create chimeric mice. After 8 weeks’ reconstitution, chimeras were used to be immunized as indicated.

### Binding antibody response against SARS-CoV-2

The general ELISA method has been previously described ^3^. Briefly, recombinant SARS-CoV-2 proteins including RBD (Sino Biological, 40592-V08H), spike S1 D614G (S1) (Sino Biological, 40591-V08H3), or nucleocapsid protein (N) (GenScript, Z03488) were precoated to 96-well plates overnight at 4°C. The next day, plates were washed with wash buffer (0.05% Tween 20 in PBS) and then blocked with assay dilution buffer (BioLegend) for 1 hour at RT. Plasma or 20 x concentrated BAL from human individuals were diluted in assay dilution buffer starting at 1:5; nasal wash was undiluted, and then serially diluted by a factor of 5. Plasma from mice was diluted generally starting at 1:500 dilution or as indicated and then serially diluted by a factor of 5. BAL from mice after SARS-CoV-2 infection was diluted starting at 1:5 dilution; BAL from mice after mucosal vaccine booster was diluted starting at 1:40 dilution or as indicated and then serially diluted by a factor of 5. Nasal wash BAL from mice was starting at undiluted and then serially diluted by a factor of 5. Samples were added to the plate and incubated for 2 hours at RT. After washing three times with wash buffer, secondary antibodies diluted with assay dilution buffer were added to the plate and then incubated for 1 hour at RT. Secondary antibodies, including anti-human IgG (Sigma-Aldrich, A6029), anti-human IgA (Hybridoma Reagent Laboratory, HP6123), anti-IgM (Sigma-Aldrich, A6907), anti-mouse IgG (SouthernBiotech, 1030-05), anti-mouse IgA (SouthernBiotech, 1040-05), and anti-mouse IgM (SouthernBiotech, 1020-05) were diluted as respectively indicated. Plates were washed three times and then developed with a 1:1 mixture of TMB substrate A and B (BioLegend) for 10 min at RT. Sulfuric acid (2 M) was used as the stop buffer. Plates were read immediately on a microplate reader (Molecular Devices) at 450 nm with SoftMax Pro Software. The optical density value at 1:5 dilution for human plasma and BAL (except for plasma IgG at 1:25); for mice, generally 1:500 dilution for plasma IgA, 1:2500 dilution for plasma IgG, 1: 40 dilution for BAL IgA, 1:200 dilution for BAL IgG, original concentration for nasal wash, was displayed, respectively. In intranasal blockade or depletion, 1:1000 dilution for BAL was displayed. For supernatant from *in vitro* experiments, 1:2 dilution was displayed.

### Neutralizing antibody response against SARS-CoV-2

Pseudovirus neutralization assays were performed as previously reported ^3^. Briefly, in a 96-well format, plasma, BAL or nasal wash was diluted starting at a 1:40, 1:20, or 1:20 dilution, respectively, and then serially diluted by a factor of 4. The pseudoviruses, including D614G, and Omicron BA.1.1, were incubated with plasma or BAL for 1 hour at 37°C, followed by infection of 2 × 10^4^ preseeded human embryonic kidney (HEK) 293T–ACE2 cells (BEI Resources, NR- 52511) on a 96-well polystyrene tissue culture plate. Gaussia luciferase activity in cell culture media was assayed 48 and 72 hours after infection. Note that, to ensure valid comparisons between SARS-CoV-2 variants, we used equivalent amounts of pseudovirus on the basis of the predetermined virus titers, and samples of different variants were loaded side by side in each plate. NT50 for each sample was determined by nonlinear regression with the least squares fit in GraphPad Prism 5 (GraphPad Software).

### Bulk RNA-sequencing and analysis

After mRNA-S plus Ad5-S immunization, CD4^+^ T cells in the BAL, lung parenchyma, and draining lymph nodes in *IL-21-VFP* reporter mice were enriched with negative selection by using biotin-labeled CD8a, TER-119, and B220 antibody and streptavidin magnetic beads (Miltenyi Biotec) before surface markers staining. Live CD4^+^CD44^+^VFP^+^ and CD4^+^CD44^+^VFP^-^ cells were sorted by Sony MA900 Cell Sorter and then resuspended in Trizol for RNA extraction. Two replicates of each cell subset were obtained. RNA quality was assessed and sequenced using Illumina 2 x 150 platform provided by Admera Health. cDNA libraries construction, sequencing alignment to the mouse reference genome through services were provided by Admera Health.

Differentially expressed genes between each group were identified using DESeq2. For functional analysis, Gene Set Enrichment Analysis was performed to identify enriched gene sets from MSigDB.

### Single-cell RNA sequencing

For library preparation, BAL cells or sorted BAL CD4^+^ T cells from immunized mice in each group were pooled together and each group was hashtagged using BioLegend TotalSeq™-A antibodies according to the provided protocol. The library of hashtagged pooled groups was then constructed using the 10x Genomics Chromium Next GEM Single Cell 3’ Reagent Kit with the v3.1 (Dual Index) protocol. Sequencing was performed using the NovaSeq X Plus platform, a service provided by Admera Health. The sequenced data were aligned using Cell Ranger. The curated data then underwent demultiplexing, quality assurance, filtering, differential gene expression analysis, and cluster identification, all conducted using the standard Seurat protocol with Seurat (v 5.1.0) in R (R-4.3.3). Downstream gene set enrichment analysis was performed using clusterProfiler (4.10.1) package, with gene sets sourced from MSigDB.

### Statistical analysis

Quantitative data are presented as means ± SEM. Statistical tests are indicated in the corresponding figure legends. Two-tailed Student’s t test (two-tailed, unequal variance) was used to determine statistical significance with Prism software (GraphPad) for two-group comparison. For multiple groups, analysis of variance (ANOVA) corrected for multiple comparisons was used when appropriate (GraphPad). Simple linear regression with 95% confidence band of the best-fit line was used for correlation analysis (GraphPad).

## Acknowledgements

We sincerely thank the study participants for their contribution to this work. We thank the NIH tetramer core facility for the T cell tetramers. We thank BEI Resources for the SARS-CoV-2 viruses. We thank the Flow Cytometry Core Facility, the Center for Comparative Medicine and the Genome Analysis Technology Core at the University of Virginia for support. The Sony MA900 Cell Sorter was funded through the NIH S10 instrument program #1S10OD028518. Schematics in the manuscript were created with BioRender.com.

The study was in part supported by the US National Institutes of Health grants AI147394, AG069264, AI112844, HL170961, and AI154598 to J.S, AI176171 to W.G.T. and J.S.; F31HL170746 and T32AI007496 to H.N; T32CA009109 to S.P.Y. American Lung Association Catalyst Award to I.S.C.; U54CA260582 to S.L.L; R01AI168684 to G. Z.; S.L.L was also supported by a fund provided by an anonymous private donor to The Ohio State University. The content is solely the responsibility of the authors and does not necessarily represent the official views of the NIH.

## Author contribution

J.T. and J.S. conceived the overall project. J.T., A.S.C., S.L.L, W.G.T. and J.S. designed the experimental strategy and analyzed data. J.T. performed majority of the experiments, A.S.C. performed the bioinformatic analysis. P.Q., P.L., Y.L., and S.L.L. performed neutralization assays. F.Z, Hui. H, T.K., H.K., M.V., Z.L., N.Z., J.J.T., G.Z and Haitao. H provided critical reagents. All other authors assisted experiments, analyzed data, or contributed critical insights to the study. J.T. and J.S. wrote the original draft. All authors read, edited, and approved the final manuscript.

## Extended Data Figure legends

**Extended Data Fig.1:**
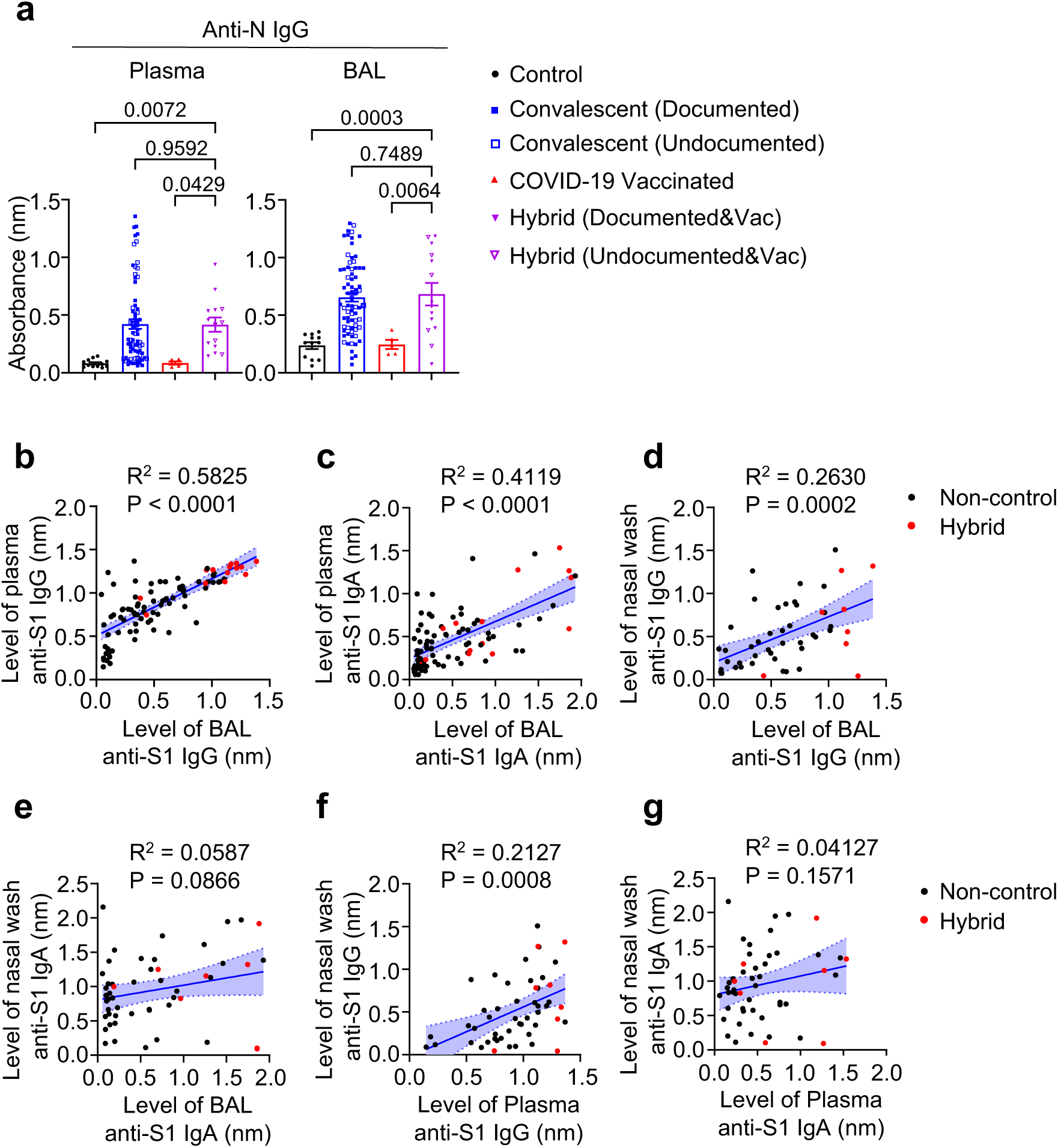
Systemic and respiratory mucosal Ab responses in individuals with SARS-CoV-2 vaccination and infection. **a,** Levels of SARS-CoV-2 N binding IgG in plasma and BAL of COVID-19 naïve individuals (n = 12), COVID-19 convalescents (n = 75), COVID-19 vaccinated individuals (n = 5), vaccinated plus infected individuals (n = 14). Individuals with undocumented COVID-19 infection are indicated as hollow symbols in groups of COVID-19 convalescents and COVID-19 vaccinated plus infected. **b-g**, Correlations between BAL IgG and plasma IgG **(b)**, BAL IgA and plasma IgA **(c)**, BAL IgG and nasal wash IgG **(d)**, BAL IgA and nasal wash IgA **(e)**, plasma IgG and nasal wash IgG **(f)**, plasma IgA and nasal wash IgA **(g)**. Individuals with hybrid immunity are indicated as red symbols. Enrolled donors’ demographics are provided in Extended Data table 1. **a**, One-way ANOVA with multiple-comparisons test. Data presented as mean ± s.e.m. **b-g**, Simple linear regression analysis with 95% confidence band of the best-fit line.

**Extended Data Fig.2:**
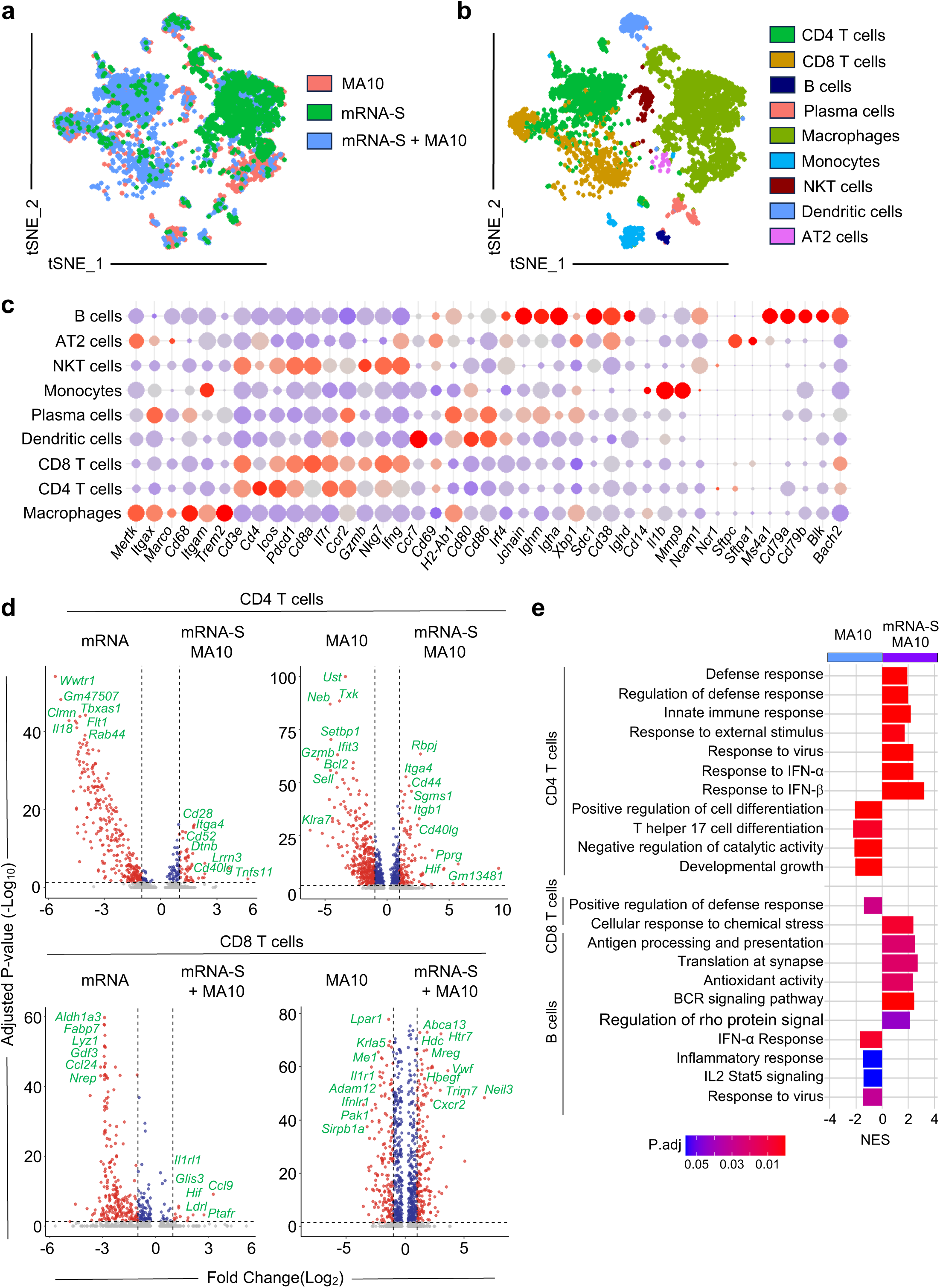
scRNA-seq analysis of hybrid immunity. **a-b,** scRNA-seq UMAP plots of BAL cells from mice at day 6 post SARS-CoV-2 MA10 infection or without infection. **c,** Heatmap of key gene expression in different clusters from scRNA-seq analysis. **d,** Volcano plots of differential expression of genes of CD4^+^ and CD8^+^ T cells in the BAL between indicated groups. **e**, Enrichment of upregulated pathways in BAL CD4^+^, CD8^+^ T and B cells from mRNA-S+ MA10 group versus MA10 group.

**Extended Data Fig.3:**
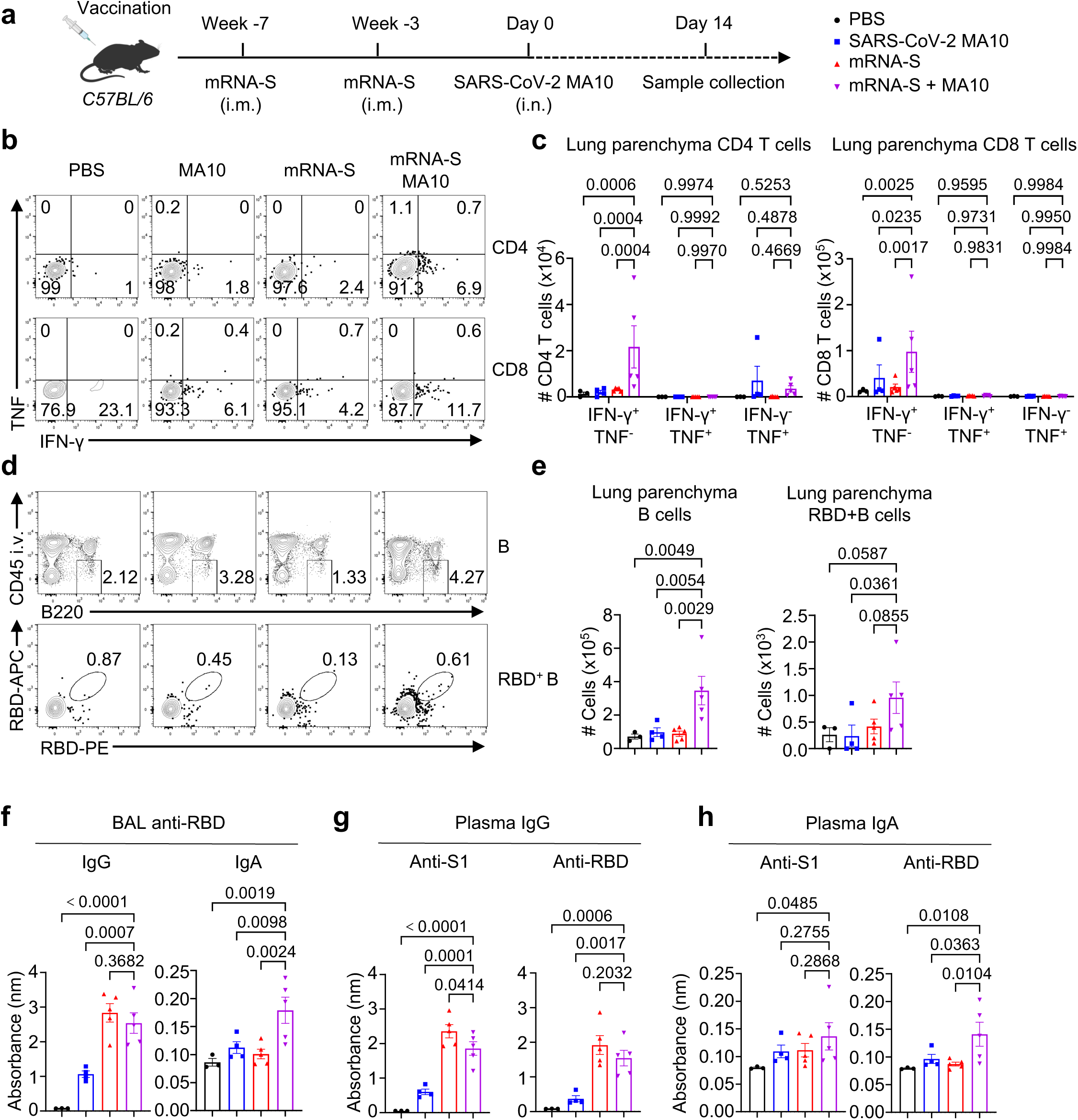
Systemic and respiratory mucosal immunity in vaccination plus SARS-CoV-2 MA10 infection model. **a,** Schematic of experimental design. C57BL/6 mice were immunized as indicated (PBS, n = 3; SARS-CoV-2 MA10, n = 4; mRNA-S, n = 5; mRNA-S + MA10, n =5). **b-e**, Represented flow cytometry plot **(b,d)** and summary **(c,e)** of SARS-CoV-2 peptide-specific CD4^+^, CD8^+^ T cell **(b,c)** and RBD-specific B cell responses **(d,e)** in lung parenchyma from different groups. **f**, SARS-CoV-2 RBD-specific IgG and IgA responses in the BAL from different groups. **g,h**, SARS-CoV-2 S1- or RBD-specific IgG **(g)** and IgA **(h)** responses in the plasma from different groups. **c**, Two-way ANOVA with multiple-comparisons test; **e-h**, one-way ANOVA with multiple-comparisons test. Data presented as mean ± s.e.m. Data are representative of two independent experiments with similar results.

**Extended Data Fig.4:**
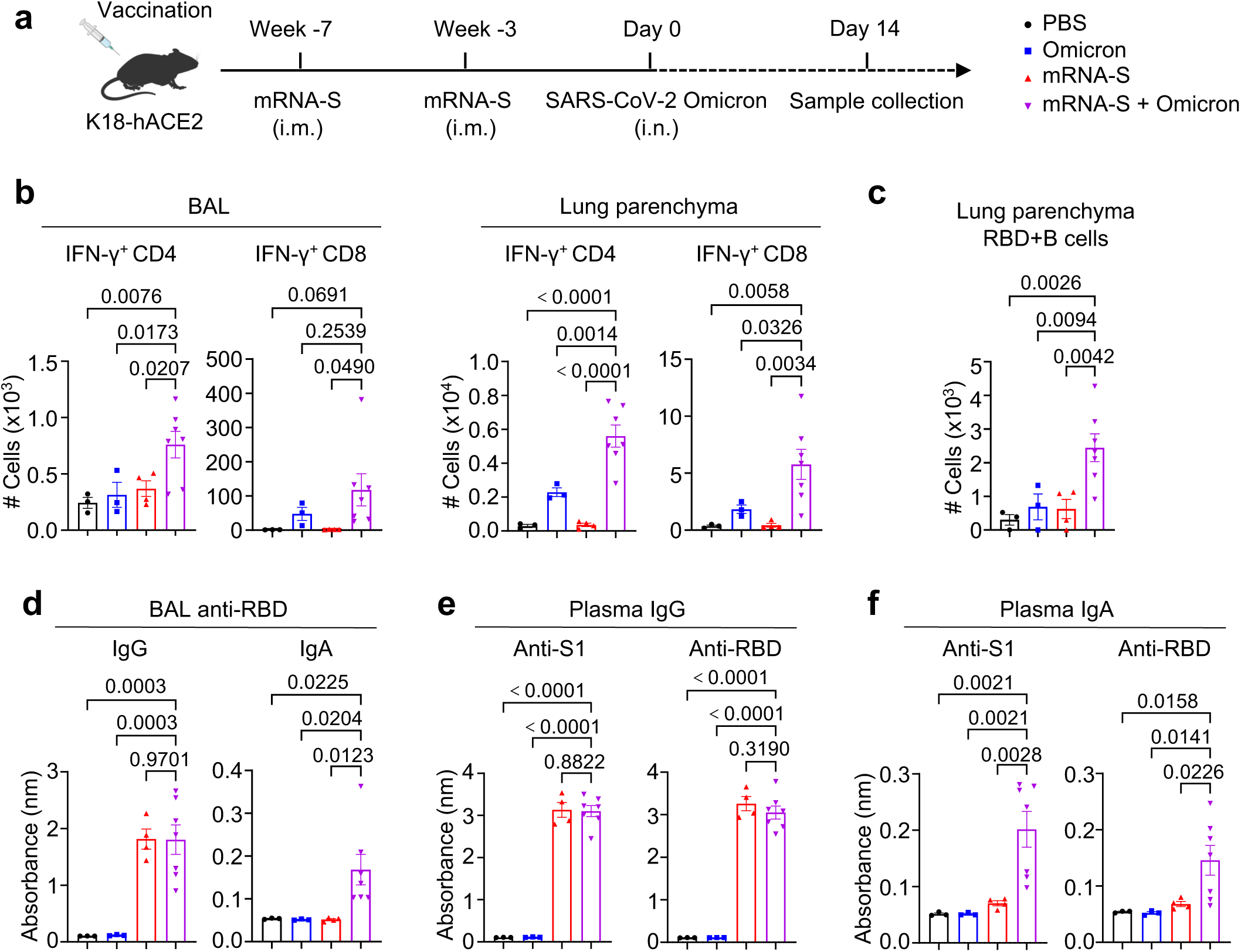
Systemic and respiratory mucosal immunity in vaccination plus SARS-CoV-2 Omicron infection model. **a,** Schematic of experimental design. K18-hACE2 mice were immunized as indicated (PBS, n = 3; SARS-CoV-2 Omicron BA.1, n = 3; mRNA-S, n = 4; mRNA-S + Omicron BA.1, n =7). **b,c**, Summary of SARS-CoV-2 peptide-specific CD4^+^, CD8^+^ T cell in the BAL and lung parenchyma **(b)** and RBD-specific B cell responses **(c)** in lung parenchyma from different groups. **d**, SARS-CoV-2 RBD-specific IgG and IgA responses in the BAL from different groups. **e,f**, SARS-CoV-2 S1- or RBD-specific IgG **(e)** and IgA **(f)** responses in the plasma from different groups. **b-f**, One-way ANOVA with multiple-comparisons test. Data presented as mean ± s.e.m. Data are representative of two independent experiments with similar results.

**Extended Data Fig.5:**
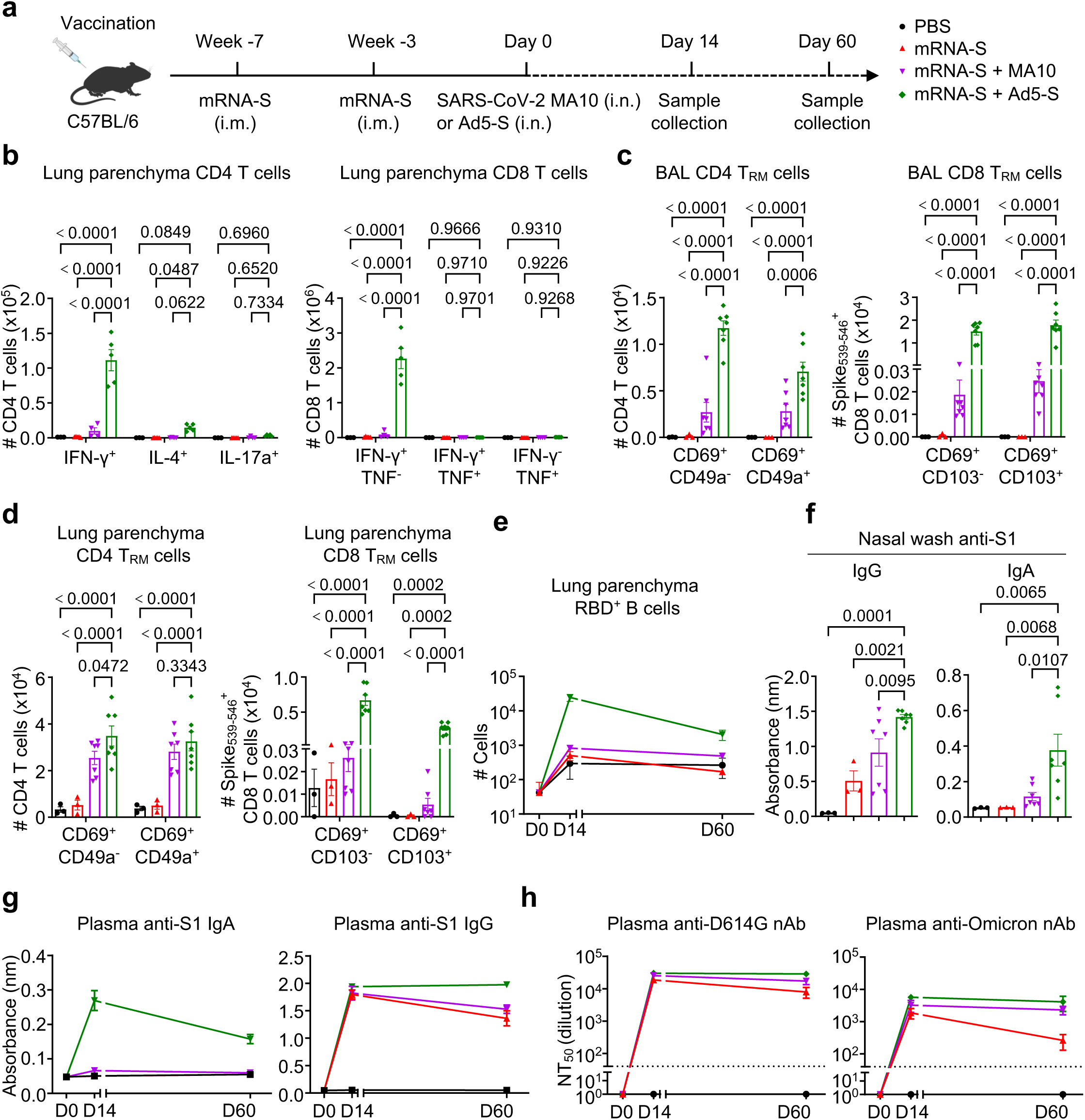
Comparative analyses of the mucosal immunity after vaccination plus SARS-CoV-2 MA10 infection, and mucosal booster vaccination. **a,** Schematic of experimental design. C57BL/6 mice were immunized as indicated (Day 0, n = 3; Day 14, PBS, n = 3; mRNA-S, n = 5; mRNA-S + MA10, n =4; mRNA-S + Ad5-S, n =5; Day 60, PBS, n = 3; mRNA-S, n = 3; mRNA-S + MA10, n =7; mRNA-S + Ad5-S, n =7). **b**, Summary of SARS-CoV-2 peptide-specific CD4^+^ T subsets, CD8^+^ T cell at day 14. **c,d**, Summary of CD4 and CD8 T_RM_ cells in the BAL **(c)** and lung parenchyma **(d)** at day 60. **e**, SARS-CoV-2 RBD-specific B cell dynamic response in lung parenchyma. **f**, SARS-CoV-2 S1-specific IgA and IgG responses in the nasal wash from different groups. **g**, SARS-CoV-2 S1-specific IgA and IgG dynamic responses in the plasma from different groups. **h**, SARS-CoV-2-specific neutralizing Ab response against D614G and Omicron BA.1 in the plasma from different groups. **b-d**, Two-way ANOVA with multiple-comparisons test; **f**, one-way ANOVA with multiple-comparisons test. Data presented as mean ± s.e.m. Data are representative of two independent experiments with similar results.

**Extended Data Fig.6:**
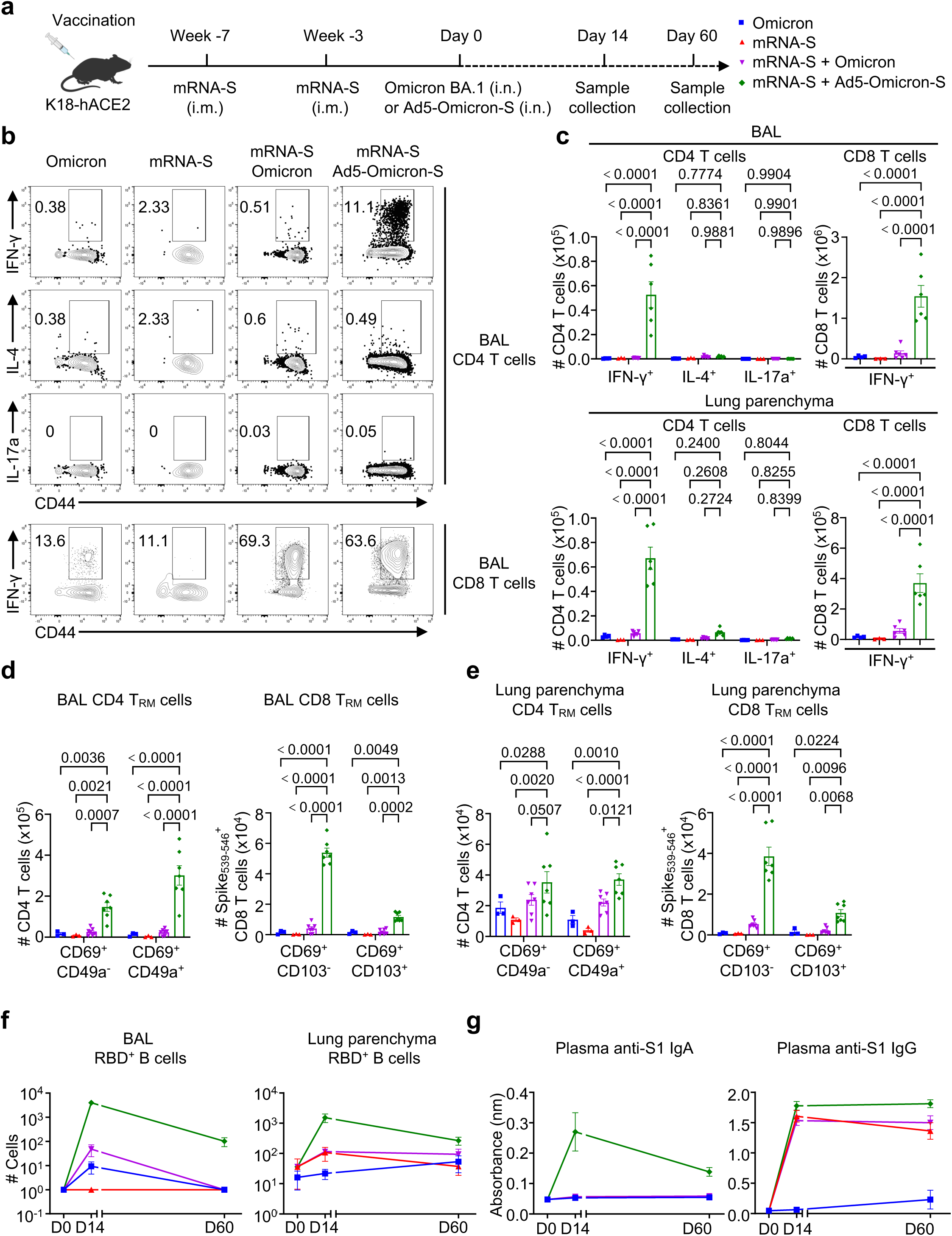
Comparative analyses of the mucosal immunity after vaccination plus SARS-CoV-2 Omicron infection, and mucosal booster vaccination. **a,** Schematic of experimental design. Schematic of experimental design. K18-hACE2 mice were immunized as indicated (Day 0, n = 3; Day 14, Omicron BA.1, n = 4; mRNA-S, n = 3; mRNA-S + Omicron BA.1, n =6; mRNA-S + Ad5-Omicron-S, n =6; Day 60, Omicron BA.1, n = 3; mRNA-S, n = 3; mRNA-S + Omicron BA.1, n =7; mRNA-S + Ad5-Omicron-S (BA.1), n =7). **b,c**, Represented flow cytometry plot **(b)** and summary **(c)** of SARS-CoV-2 peptide-specific CD4^+^ T subsets, CD8^+^ T cell at day 14. **d,e**, Summary of CD4 and CD8 T_RM_ cells in the BAL **(d)** and lung parenchyma **(e)** at day 60. **f**, SARS-CoV-2 RBD-specific B cell dynamic response in the BAL and lung parenchyma. **g**, SARS-CoV-2 S1-specific IgA and IgG dynamic responses in the plasma from different groups. **c-e**, Two-way ANOVA with multiple-comparisons test. Data presented as mean ± s.e.m. Data are representative of two independent experiments with similar results.

**Extended Data Fig.7:**
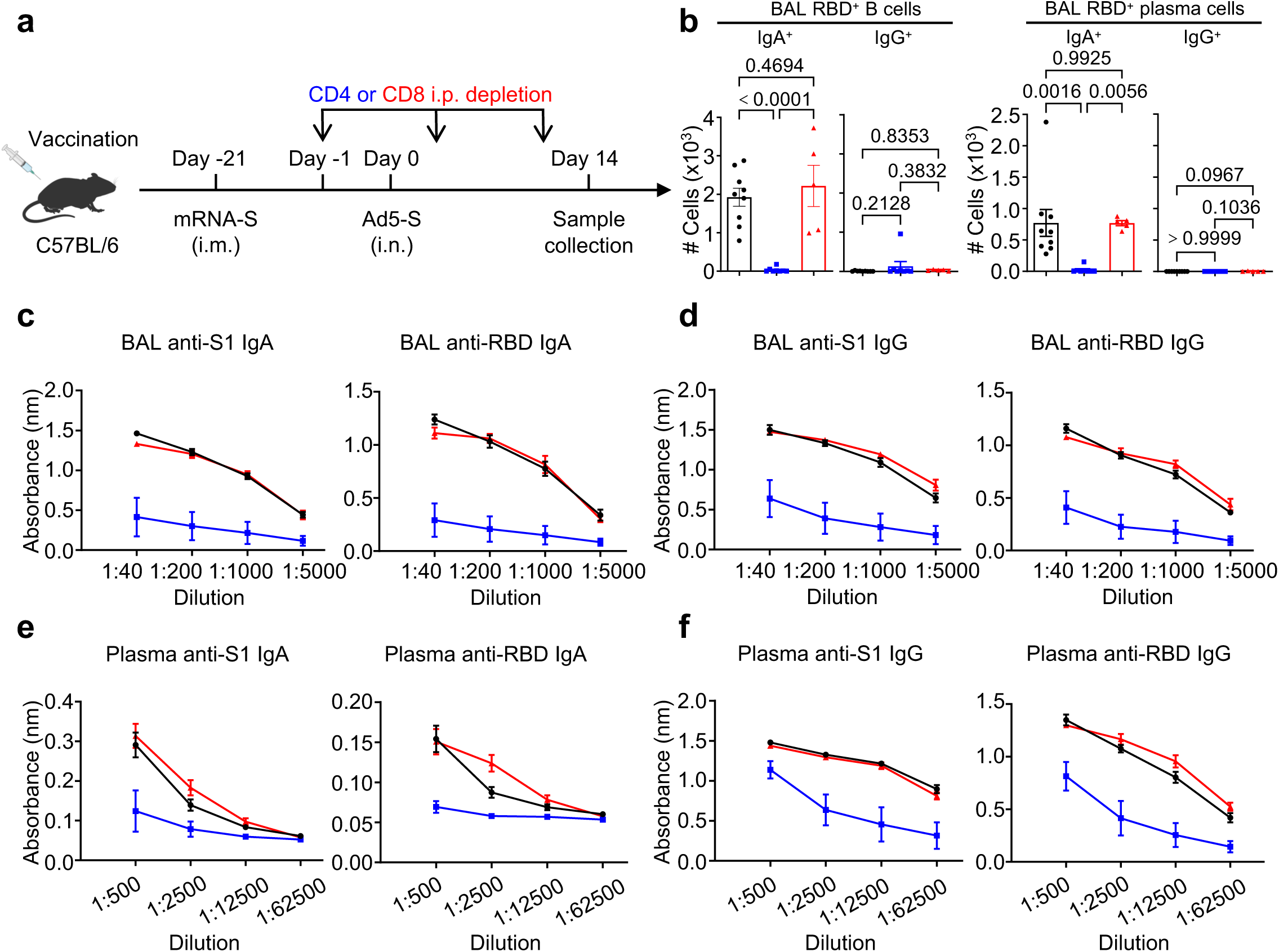
CD4^+^, but not CD8^+^, T cell help is required for mucosal IgA responses during mucosal booster. **a-f,** C57BL/6 mice were immunized as indicated (ctrl, n = 9; CD4 i.p. depletion, n = 8; CD8 i.p. depletion, n = 5). Schematic of experimental design **(a)**; SARS-CoV-2 RBD-specific IgA or IgG positive B cell and plasma cell responses **(b)** in the BAL. SARS-CoV-2 S1- or RBD-specific IgA **(c,e)** and IgG **(d,f)** responses in the BAL **(c,d)** and plasma **(e,f)**. **b**, One-way ANOVA with multiple-comparisons test. Data presented as mean ± s.e.m. Data are pooled from two independent experiments.

**Extended Data Fig.8:**
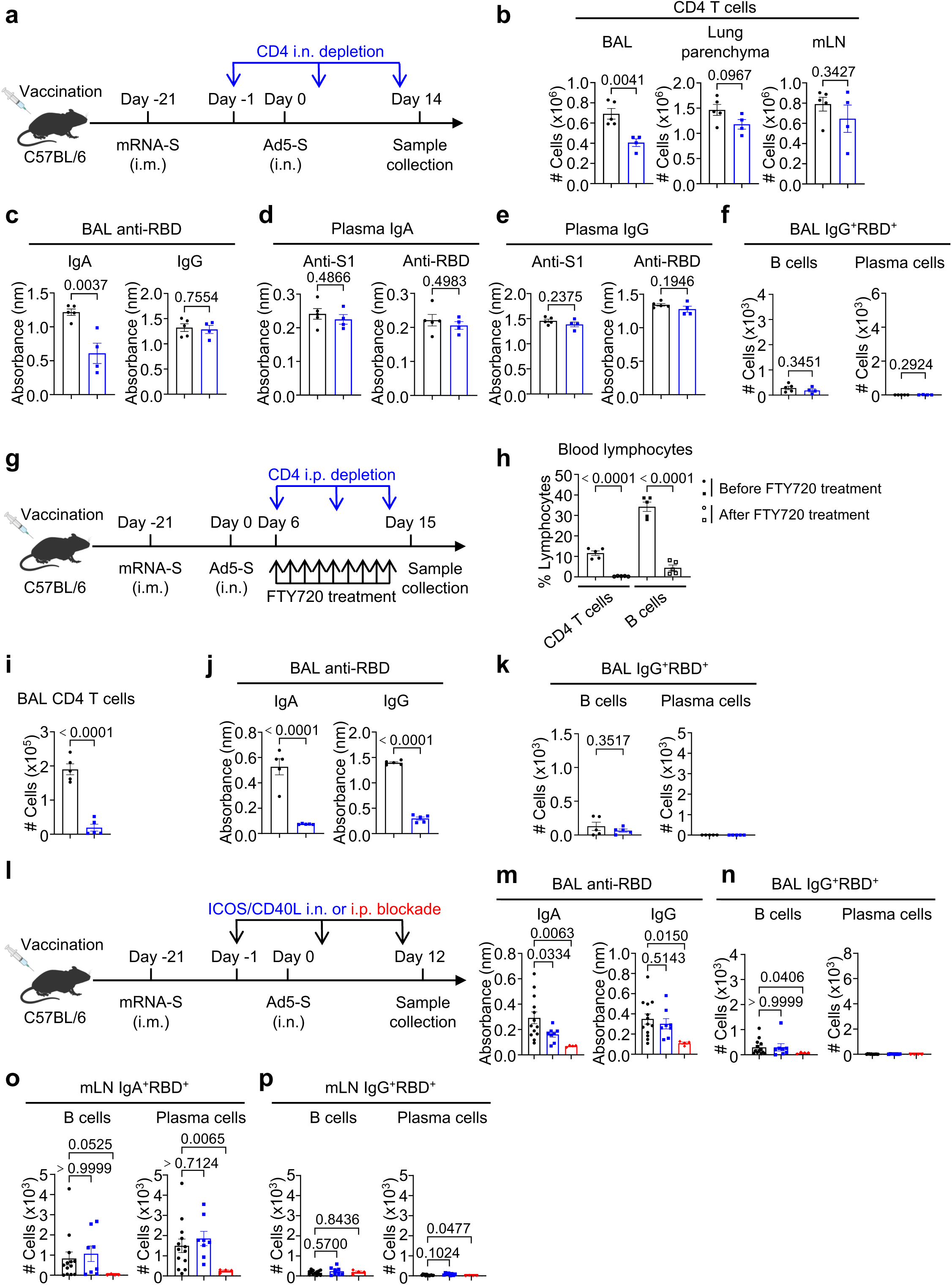
Optimal mucosal IgA responses are supported by T-B interactions *in situ*. **a-f,** C57BL/6 mice were immunized as indicated (Ctrl, n = 5; CD4 i.n. depletion, n = 4). Schematic of experimental design **(a)**; CD4^+^ T cell responses in the BAL, lung parenchyma and draining lymph nodes **(b)**; SARS-CoV-2 RBD-specific IgA and IgG responses in the BAL **(c);** SARS-CoV-2 S1- or RBD-specific IgA **(d)** and IgG **(e)** responses in the plasma and SARS-CoV- 2 RBD-specific IgG positive B cell and plasma cell responses in the BAL **(f)**. **g-k**, C57BL/6 mice were immunized as indicated (FTY720 treated ctrl, n = 5; FTY720 treatment plus CD4 i.p. depletion, n =4). Schematic of experimental design **(g)**; Percentage of CD4^+^ T and B cells in the blood before or after FTY720 treatment **(h)**; CD4^+^ T cell response in the BAL **(i)**; SARS-CoV-2 RBD-specific IgA and IgG responses **(j)** and SARS-CoV-2 RBD-specific IgG positive B cell and plasma cell responses **(k)** in the BAL. **l-p**, C57BL/6 mice were immunized as indicated (Ctrl, n = 13; ICOSL and CD40L i.n. blockade, n =8; ICOSL and CD40L i.p. blockade, n =4). Schematic of experimental design **(l)**; SARS-CoV-2 RBD-specific IgA and IgG responses **(m)**; SARS-CoV-2 RBD-specific IgG positive B cell and plasma cell responses **(n)** in the BAL; SARS-CoV-2 RBD- specific IgA **(o)** or IgG **(p)** positive B cell and plasma cell responses in the draining lymph nodes. **b-f, i-k**, Unpaired two-sided t-test; **h**, two-way ANOVA with multiple-comparisons test; **m-p**, one-way ANOVA with multiple-comparisons test. Data presented as mean ± s.e.m. **a-k**, Data are representative of two independent experiments with similar results. **l-p**, Data are pooled from two independent experiments.

**Extended Data Fig.9:**
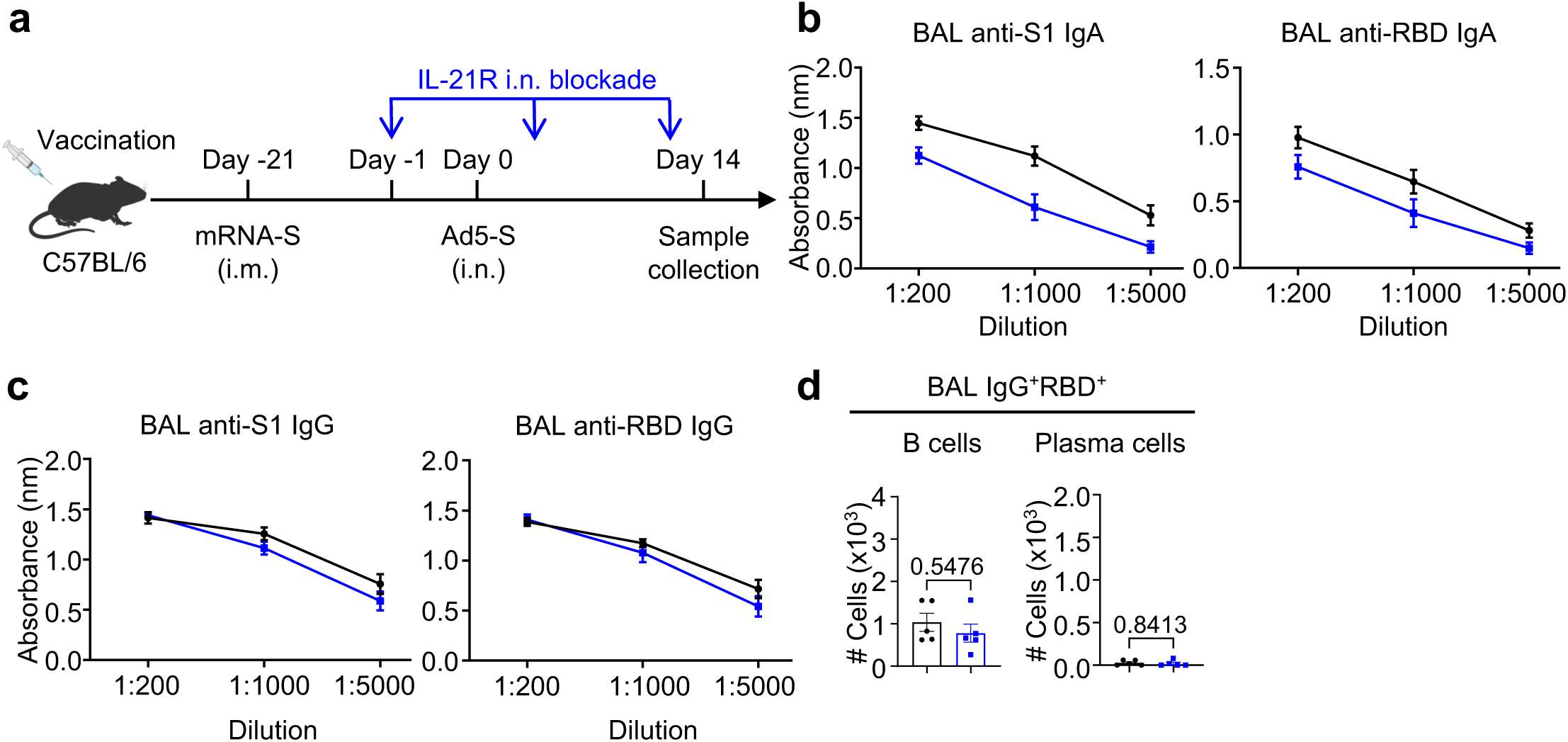
Optimal mucosal IgA responses are supported by local IL-21 signaling. **a-d,** C57BL/6 mice were immunized as indicated (Ctrl, n = 5; IL-21R i.n. blockade, n = 5). Schematic of experimental design **(a)**; SARS-CoV-2 S1- or RBD-specific IgA **(b)** and IgG **(c)** antibody responses and SARS-CoV-2 RBD-specific IgG positive B cell and plasma cell responses **(d)** in the BAL. **d**, Unpaired two-sided t-test. Data presented as mean ± s.e.m. Data are representative of two independent experiments with similar results.

**Extended Data Fig.10:**
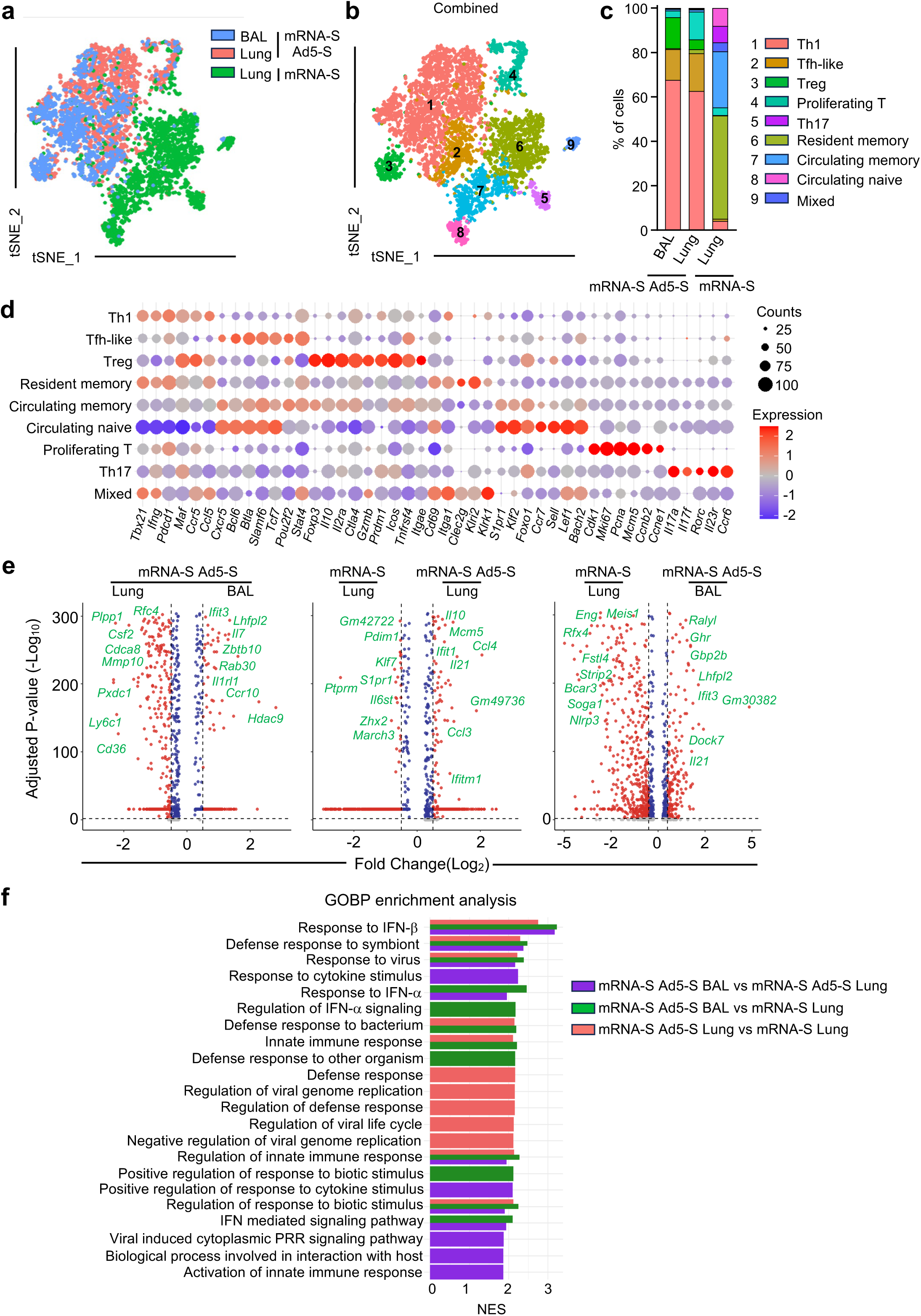
ScRNA-seq analysis of CD4 T cells after mRNA vaccination alone or mucosal booster. **a-b**, scRNA-seq UMAP plots **(a,b)** with percentage of different clusters **(c)** of BAL cells from mice at day 6 post Ad5-S booster and lung parenchyma CD4 T cells from mice without Ad5-S booster (mRNA + Ad5-S, n = 3; mRNA, n = 3). **d**, Heatmap of key gene expression in different clusters from scRNA-seq analysis. **e,** volcano plots of differential expression of genes of CD4^+^ T cells between indicated groups. **f**, GOBP enrichment pathways from scRNA-seq analysis.

**Extended Data Fig.11:**
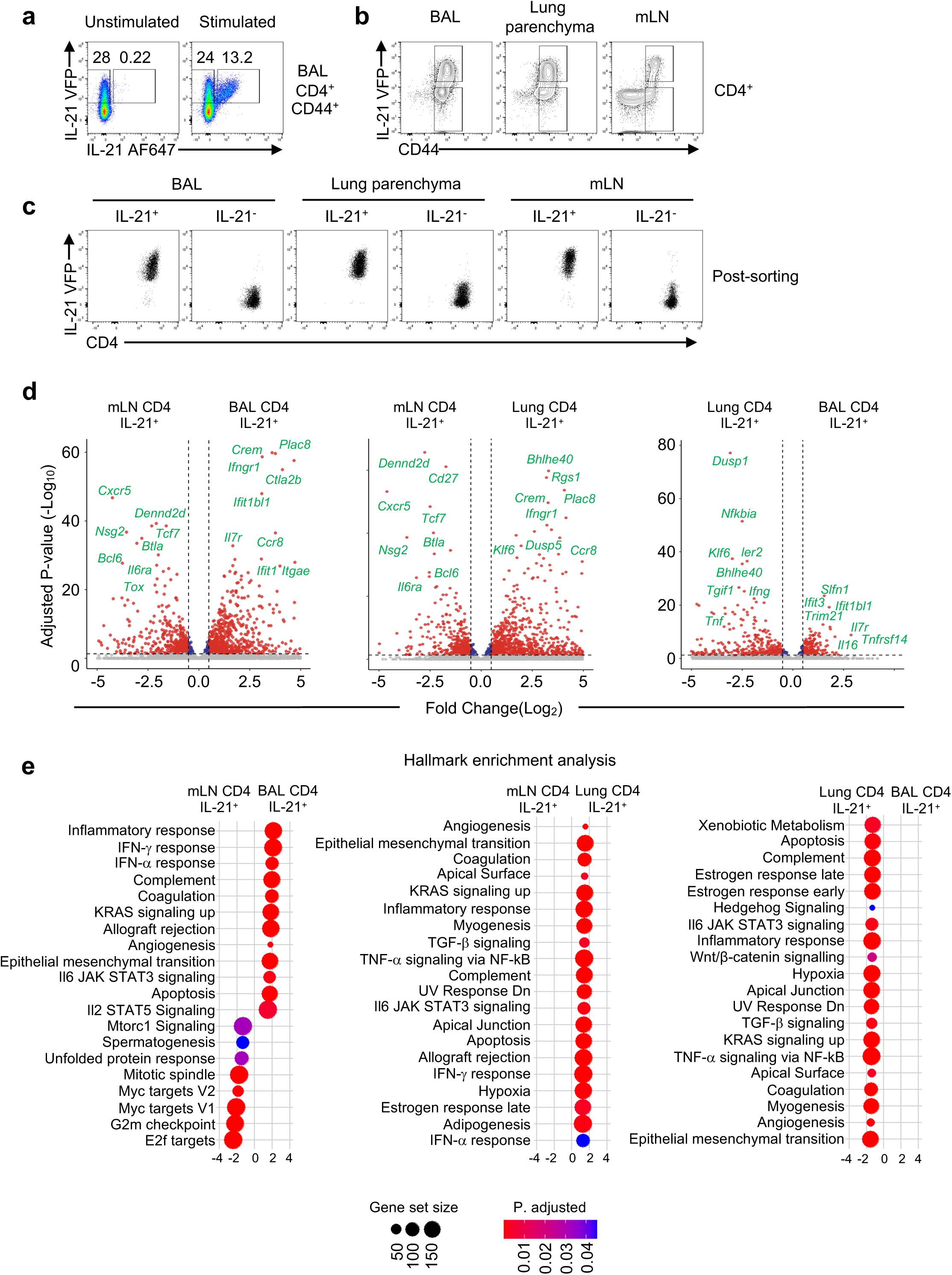
Molecular signatures of respiratory IL-21^+^ CD4 T cells after mucosal booster. **a**, Represented flow cytometry plot of IL-21 intracellular staining and IL-21-VFP in BAL CD4^+^CD44^+^ T cells from mice with Ad5-S booster. **b,c**, Sorting strategy **(b)** and post-sorting check **(c)** of IL-21^+^ and IL-21^-^ CD4^+^ T cell in the BAL, lung parenchyma and draining lymph nodes. **d,e**, Bulk RNA-seq analysis of IL-21^+^ and IL-21^-^ CD4^+^ T cells in BAL, lung parenchyma and mLN from IL-21-VFP reporter mice at day 6 post Ad5-S booster (n = 5). Volcano plots of differential expression of genes **(c)** and Hallmark enrichment analysis **(d)** between indicated groups.

**Extended Data Fig.12:**
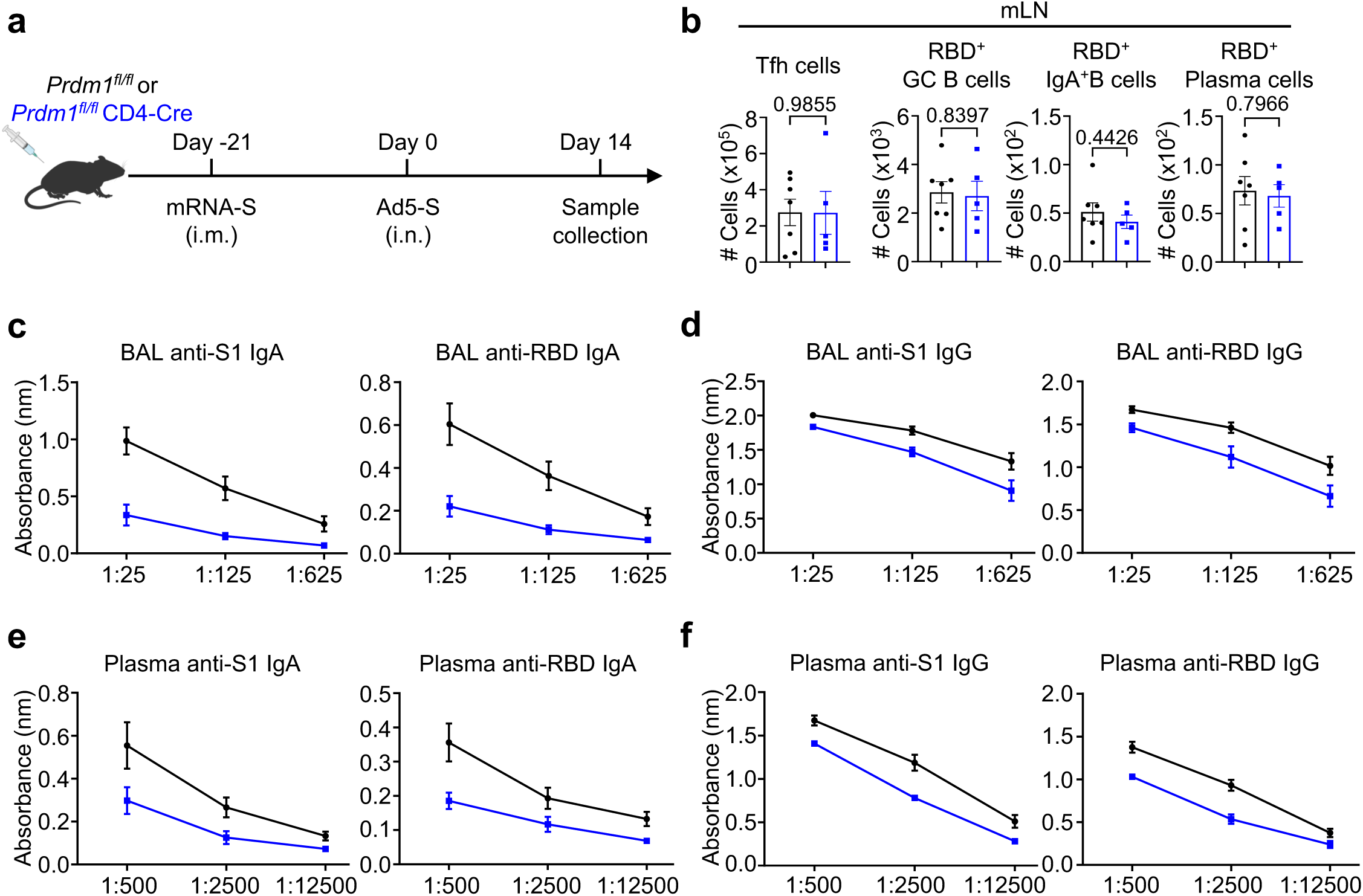
Optimal mucosal IgA responses are supported by Blimp-1^+^ Th1 effector cells. **a-f,** *Prdm1^fl/fl^* and *Prdm1^fl/fl^* CD4-Cre mice were immunized with mRNA-S and Ad5-S as previously (*Prdm1^fl/fl^*, n = 7; *Prdm1^fl/fl^* CD4-Cre, n = 5). Schematic of experimental design **(a)**; Summary of Tfh cells (CXCR5^+^PD-1^+^), SARS-CoV-2 RBD-specific GC B cell, IgA^+^ B cell and plasma cell responses in the draining lymph nodes **(b)**; SARS-CoV-2 S1- or RBD-specific IgA **(c,e)** and IgG **(d,f)** antibody responses in the BAL **(c,d)** and plasma **(e,f)**. **b**, Unpaired two-sided t-test; Data presented as mean ± s.e.m.

**Extended Data Fig.13:**
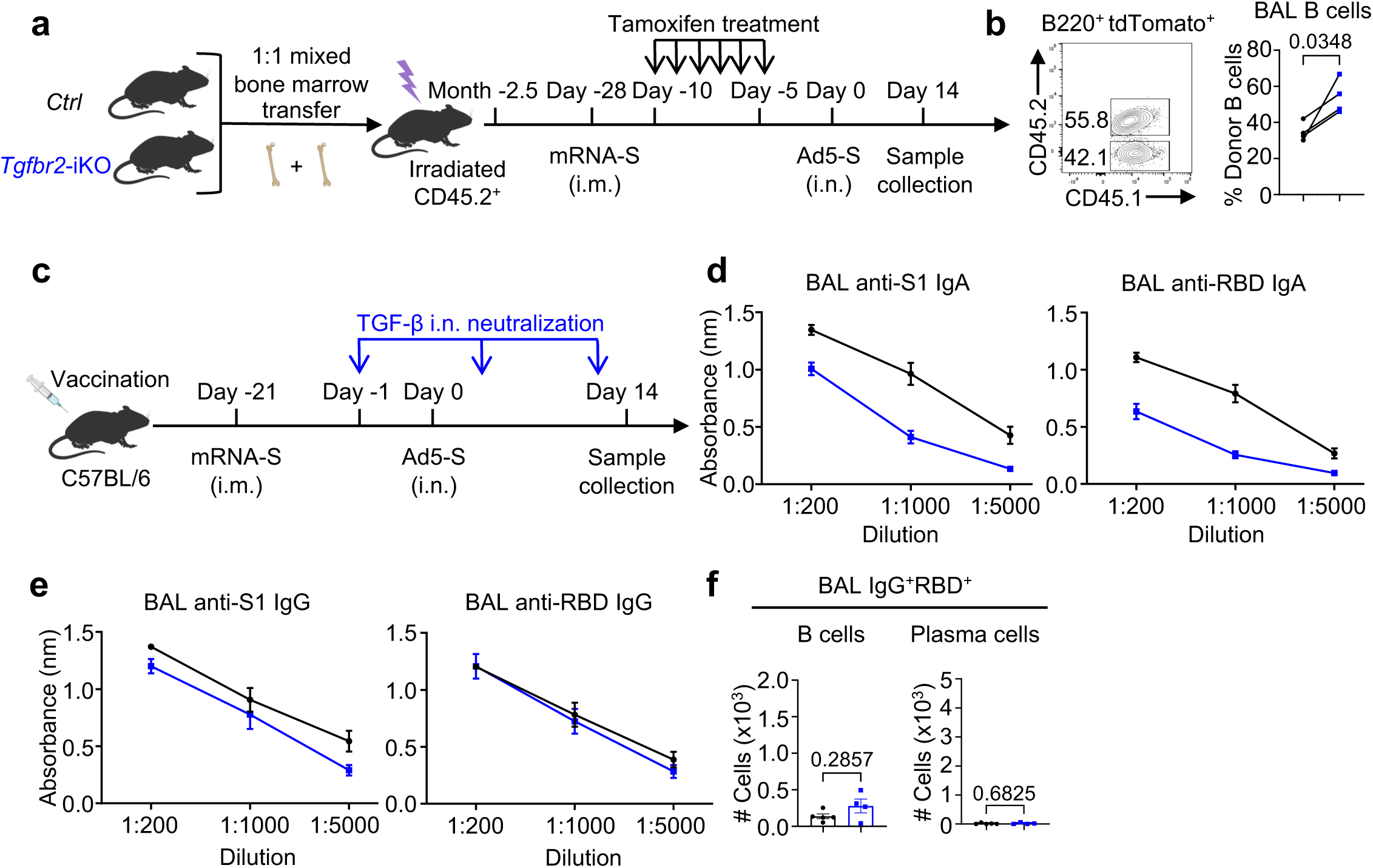
Mucosal IgA^+^ B cell responses require B cell-intrinsic TGF-β signaling. **a,b**, Mixed CD45.1^+^R26-CreERT2-tdTomato (Ctrl) and CD45.1^+^CD45.2^+^R26-CreERT2- tdTomato *Tgfbr2^fl/fl^ (*Tgfbr2-iKO) BM (1:1) chimeric mice were immunized and treated as indicated (n = 4). Schematic of experimental design **(a)**; Represented flow cytometry plot and summary of B cell percentage of different compartments **(b)**. **c-f**, C57BL/6 mice were immunized as indicated (Ctrl, n = 5; TGF-β i.n. blockade, n = 4). Schematic of experimental design **(c)**; SARS-CoV-2 S1- or RBD-specific IgA **(d)** and IgG **(e)** antibody responses and SARS-CoV-2 RBD-specific IgG positive B cell and plasma cell responses **(f)** in the BAL. **b**, Paired two-sided t-test. Data presented as mean ± s.e.m. Data are representative of two independent experiments with similar results.

**Extended Data Fig.14:**
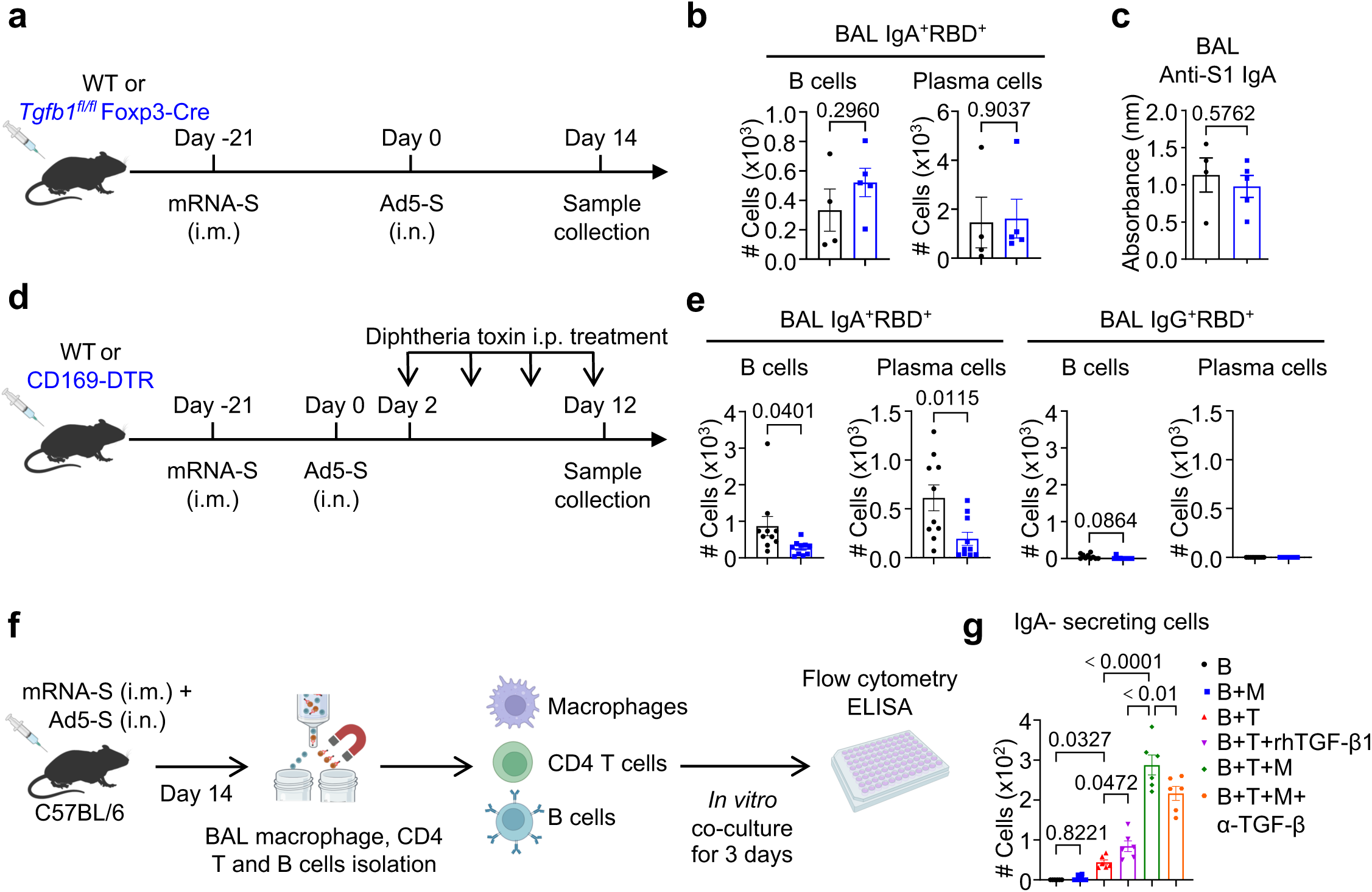
Lung macrophages and TGF-β assist optimal mucosal IgA production. **a-c**, WT and *Tgfb1^fl/fl^* Foxp3-Cre mice were immunized with mRNA-S and Ad5-S as previously (WT, n = 4; *Tgfb1^fl/fl^* Foxp3-Cre, n = 5). Schematic of experimental design **(a)**; SARS-CoV-2 RBD-specific IgA positive B cell and plasma cell responses **(b)** and SARS-CoV-2 S1-specific IgA antibody responses **(c)** in the BAL. **d,e**, WT and CD169-DTR mice were immunized with mRNA-S and Ad5-S as previously (WT, n = 10; CD169-DTR, n = 10). Schematic of experimental design **(d)**; SARS-CoV-2 RBD-specific IgA or IgG positive B cell and plasma cell responses **(e)**. **f-g**, C57BL/6 mice were immunized with mRNA-S and Ad5-S as previously (n = 5). BAL CD4^+^ T, B cells and macrophages were isolated at day 14 for *in vitro* coculture for 3 days. Schematic of experimental design **(f)** and IgA secreting cells number in different groups **(g)**. **b,c,e**, Unpaired two-sided t-test; **g**, one-way ANOVA with multiple-comparisons test. Data presented as mean ± s.e.m. **d,e**, Data are pooled from two independent experiments; **f,g**, data are representative of two independent experiments with similar results.

**Extended Data Fig.15:**
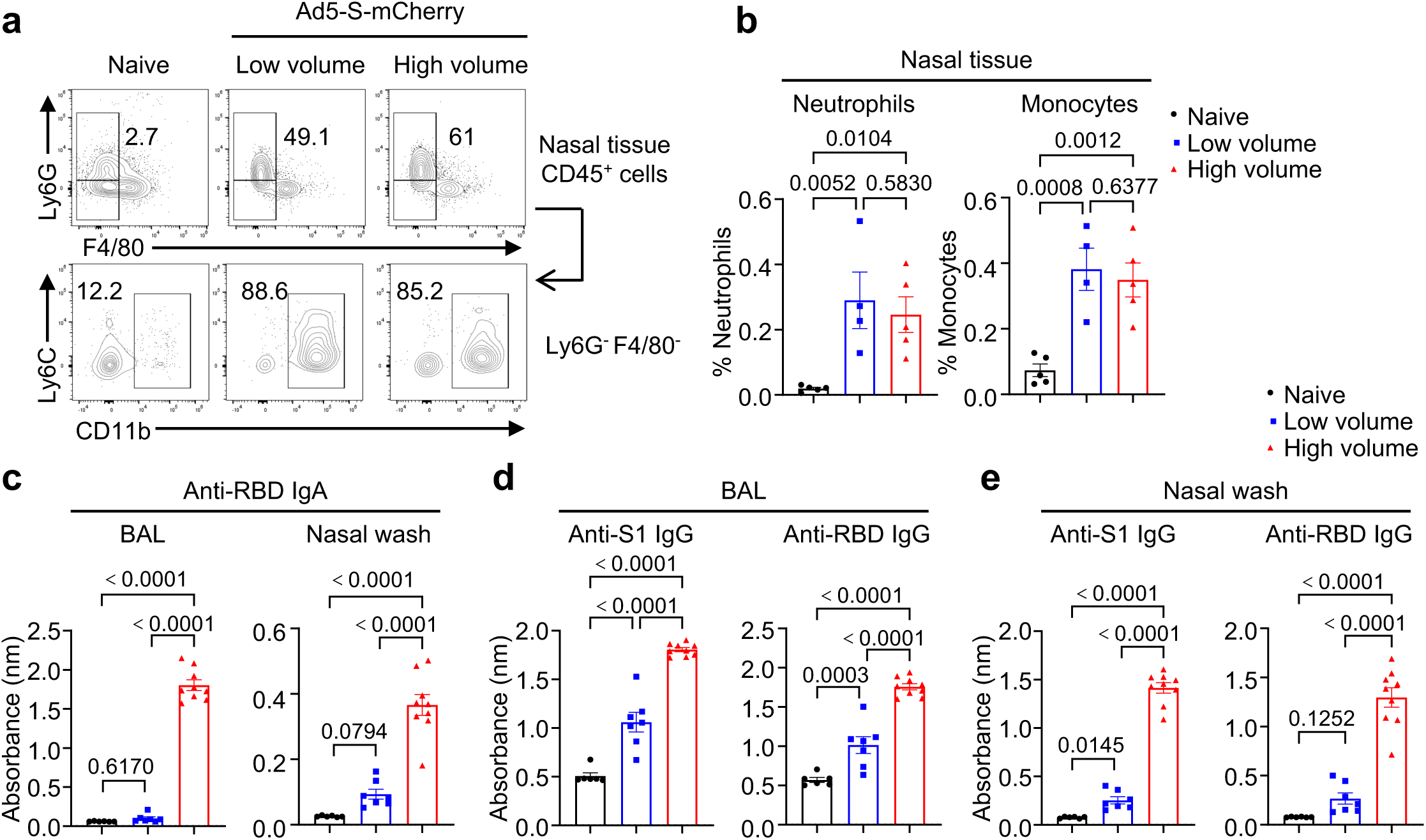
Targeting lung macrophages promotes optimal mucosal IgA production. **a,b**, C57BL/6 mice were immunized with one dose of mRNA-S plus Ad5-S-mCherry (Naïve, n = 5; low volume, n = 4; high volume, n = 5). Represented flow cytometry plot **(a)** and summary **(b)** of neutrophiles and monocytes in the nasal tissue at day 2.5 post Ad5-S. **c-e**, SARS-CoV-2 RBD-specific IgA responses in the BAL and nasal wash **(c)** and S1- or RBD-specific IgG responses in the BAL **(d)** and nasal wash **(e)** from mice immunized with mRNA-S and Ad5-S as previously at day 14 (Naïve, n = 6; low volume, n = 7; high volume, n = 9). ). **b-e**, One-way ANOVA with multiple-comparisons test. Data presented as mean ± s.e.m. **a-e**, Data are pooled from two independent experiments.

**Extended Data Fig.16:**
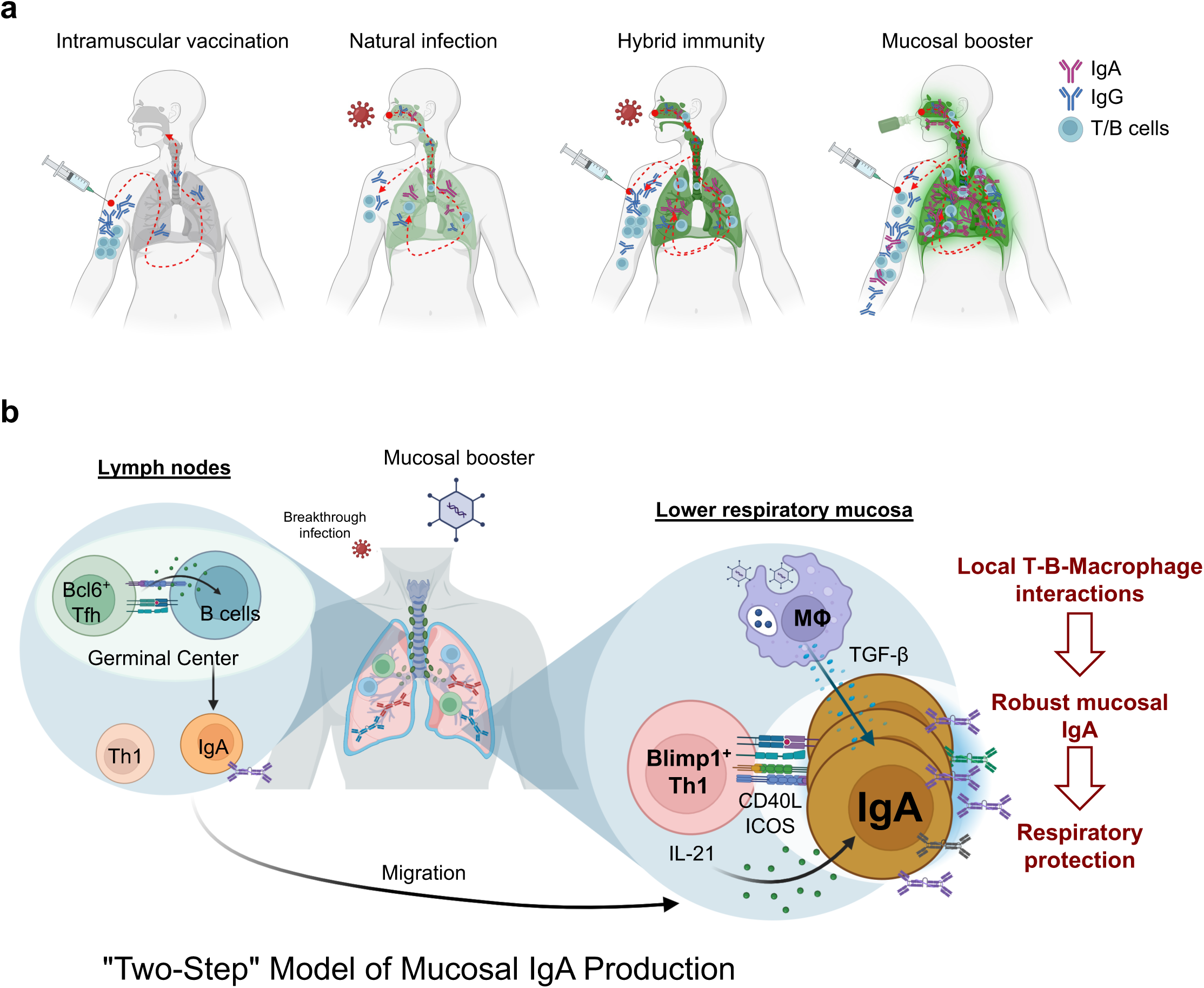
Model on the mechanism of mucosal IgA production. **a,** Current intramuscular mRNA vaccination induces relatively weak mucosal immunity, primarily through IgG antibodies. Natural SARS-CoV-2 infection generates moderate mucosal immunity. A combination of vaccination and infection results in stronger mucosal immunity. However, intramuscular vaccination followed by a mucosal booster elicits robust mucosal immunity with superior IgA responses. **b**, “Two-Step” Model of Mucosal IgA Production: The first step involves intramuscular vaccination, which initiates the development of IgA+ B cells in lymphoid organs, facilitated by Bcl6^+^ Tfh cells. In the second step, lymphocytes migrate from lymphoid organs to the respiratory mucosa, where IL-21-producing Blimp-1^+^ Th1 cells and TGF-β-producing lung macrophages cooperatively support an optimal mucosal IgA-secreting cell response, leading to enhanced IgA production to confer optimal mucosal protection.

---

**Extended Data table 1:** Study Participants Characteristics.

|  | Uninfected/<br>Unvaccinated | Uninfected/<br>Vaccinated | Infected/<br>Unvaccinated | Infected/<br>Vaccinated |
| --- | --- | --- | --- | --- |
|  | n = 12 | n = 5 | n = 75 | n = 14 |
| Category Name | Control | Vaccine | COVID | Hybrid |
| Age (years) <sup>+</sup> | 2.7 ± 1.7 <sup>γ</sup> | 10.7 ± 5.5 | 4.6 ± 3.1 <sup>†</sup> | 7.6 ± 4.1 |
| Male sex n (%) | 10 (83) | 4 (71) | 58 (77) | 11 (80) |
| Race/ethnicity n (%) |  |  |  |  |
| White | 11 (92) | 4 (80) | 59 (79) | 9 (64) |
| Black | 1 (8) | 1 (20) | 12 (16) | 4 (29) |
| Mixed | 0 | 0 | 4 (5) | 1 (7) |
| Hispanic/Latino n (%) | 0 | 0 | 3 (4) | 0 |
| Body Mass Index (%ile) <sup>*</sup> | 93 ± 43 | 62 ± 30 | 76 ± 47 | 67 ± 49 |
| Social vulnerability <sup>α*</sup> | 0.44 ± 0.34 | 0.47 ± 0.39 | 0.34 ± 0.37 | 0.34 ± 0.25 |
| Annual median ZIP code<br>income/household (\$) <sup>*</sup> | 54,594 ±<br>25,517 | 75,945 ±<br>24,042 | 58,544 ±<br>27,887 | 62,009 ±<br>46,364 |
| Residence population<br>density (per square mile) <sup>*</sup> | 203 ± 222 | 266 ± 641 | 167 ± 326 | 102 ± 315 |
| Urban residence | 2 (17) | 1 (20) | 14 (18) | 2 (14) |
| Days post-infection <sup>*</sup> |  |  | 199 ± 292 | 172 ± 301 |
| Days post-vaccine <sup>*</sup> |  | 370 ± 541 |  | 200 ± 284 |
| Age onset symptoms <sup>*</sup> | 6 ± 18 | 12 ± 99 | 12 ± 18 | 12 ± 28 |
| Duration of symptoms<br>(months) <sup>*</sup> | 13 ± 30 | 71 ± 100 | 25 ± 42 <sup>Δ</sup> | 61 ± 70 |
| Daily ICS dose (mcg<br>fluticasone equivalents) <sup>*</sup> | 242 ± 393 | 440 ± 100 | 440 ± 440 | 308 ± 476 |
| High-dose ICS therapy | 3 (25) | 2 (40) | 22 (45) | 6 (43) |
| Anti-IL-5Ra Rx n (%) | 0 | 1 (20) | 1 (1) | 0 |
| Anti-IgE Rx n (%) | 0 | 1 (20) | 1(1) | 1 (7) |
| Co-morbid Diagnoses |  |  |  |  |
| Recurrent croup | 7 (58) | 3 (60) | 39 (52) | 9 (64) |
| GERD | 5 (42) | 3 (60) | 39 (52) | 6 (43) |
| Premature birth < 36 wks | 0 | 1 (20) | 11 (15) | 4 (29) |
| Eosinophilic esophagitis | 1 (8) | 0 | 9 (12) | 1 (7) |
| Serum total IgE (IU/ml) <sup>*</sup> | 8 ± 49 | 50 ± 123 | 36 ± 139 | 67 ± 224 |
| # positive allergen tests + | 1 ± 3 | 3 ± 5 | 3 ± 5 | 5 ± 6 |
| Any positive allergen | 9 (82) | 3 (60) | 44 (59) | 7 (50) |
| Total blood eosinophils <sup>*</sup> | 170 ± 320 | 80 ± 250 | 230 ± 320 | 260 ± 290 |
| Blood eosinophils > 300 | 17 (44) | 3 (43) | 16 (33) | 5 (45) |
| Total blood neutrophils <sup>*</sup> | 3180 ± 1630 | 2930 ± 3380 | 3045 ± 2590 | 4370 ± 3860 |
| Total blood lymphocytes <sup>*</sup> | 4135 ± 980 | 2440 ± 1140 | 3040 ± 1460 <sup>Ω</sup> | 2620 ± 1030 |
| Blood hs-CRP (mg/dl) | 0.3 ± 2.0 | 0.1 ± 0.1 | 0.3 ± 1.7 | 0.3 ± 3.0 |
| <sup>+</sup> mean ± 1 standard deviation; <sup>γ</sup> p < 0.05 compared to vaccine and hybrid, One-way ANOVA with Bonferroni correction; <sup>†</sup> p < 0.05 compared to vaccine and hybrid, <sup>*</sup> median ± interquartile range; <sup>α</sup> Centers for Disease Control Social Vulnerability Index scale 0 to 1; <sup>Ω</sup> P < 0.05 compared to control; <sup>Δ</sup> p < 0.05 compared to control and hybrid. |  |  |  |  |

**Extended Data table 2:** Lung Lavage Features and Pathogen Burden.

|  | Uninfected/<br>Unvaccinated | Uninfected/<br>Vaccinated | Infected/<br>Unvaccinated | Infected/<br>Vaccinated |
| --- | --- | --- | --- | --- |
|  | n = 12 | n = 5 | n = 75 | n = 14 |
| Category Name | Control | Vaccine | COVID | Hybrid |
| Total cell count X 10<sup>6</sup>/ml * | 0.25 ± 0.32 | 0.11 ± 0.13 | 0.10 ± 0.29 | 0.39 ± 1.60 |
| Eosinophils (%) + | 0.4 ± 1.2 | 1.0 ± 1.7 | 0.3 ± 1.4 | 1.1 ± 3.7 |
| Neutrophils (%) * | 2.0 ± 52 | 2.0 ± 13 | 4.0 ± 46 | 9.0 ± 50 |
| Lymphocytes (%) * | 0.5 ± 4.0 | 4.0 ± 3.0 | 2.0 ± 5.0 | 3.0 ± 3.0 |
| Macrophages (%) * | 79.0 ± 62.0 | 71.0 ± 28.0 | 55.5 ± 50.0 | 67.5 ± 55.0 |
| Epithelial cells (%) * | 7.5 ± 11.0 | 10.0 ± 24.0 | 6.0 ± 24.0 | 3.5 ± 17.0 |
| BAL Granulocyte Patterns |  |  |  |  |
| Differential not done | 0 | 0 | 3 (4) | 0 |
| Isolated eosinophilia | 1 (8) | 1 (20) | 6 (8) | 0 |
| Isolated neutrophilia | 3 (25) | 0 | 27 (36) | 6 (43) |
| Mixed granulocytosis | 1 (8) | 1 (20) | 3 (4) | 3 (21) |
| Pauci-granulocytic | 7 (58) | 3 (60) | 36 (48) | 5 (36) |
| Lung Lavage Pathogens |  |  |  |  |
| No pathogens n (%) | 5 (42) | 5 (100) | 40 (56) | 9 (64) |
| Rhinovirus alone n (%) | 4 (33) | 0 | 16 (22) | 1 (7) |
| Pathogenic bacteria n (%) | 1 (8) | 0 | 2 (3) | 1 (7) |
| Combination virus/bact. | 0 | 0 | 7 (10) | 2 (14) |
| Non-RV resp. viruses | 2 (17) | 0 | 7 (10) | 1 (7) |
| * median ± interquartile range; + mean ± one standard deviation. |  |  |  |  |

**Extended Data table 3:** Antibody list for flow cytometry.

| ANTIGEN | FLUORESCENCE | CLONE | CATALOGUE NUMBER | COMPANY |
| --- | --- | --- | --- | --- |
| CD103 | Brilliant Violet 421 | 2E7 | 121422 | BioLegend |
| CD103 | Brilliant Violet 605 | 2E7 | 121433 | BioLegend |
| CD11b | PE | M1/70 | 101208 | BioLegend |
| CD11c | Brilliant Violet 510 | N418 | 117338 | BioLegend |
| CD138 | Brilliant Violet 605 | 281-2 | 142531 | BioLegend |
| CD19 | Brilliant Violet 650 | 6D5 | 115541 | BioLegend |
| CD38 | Brilliant Violet 421 | 90 | 102732 | BioLegend |
| CD38 | Alexa Fluor 700 | 90 | 102742 | BioLegend |
| CD4 | Brilliant Violet 785 | GK1.5 | 100453 | BioLegend |
| CD4 | Brilliant Violet 785 | RM4-5 | 100552 | BioLegend |
| CD44 | Brilliant Violet 605 | IM7 | 103047 | BioLegend |
| CD45 | PerCP/Cyanine5.5 | 30-F11 | 103132 | BioLegend |
| CD45 | Brilliant Violet 785 | 30-F11 | 103149 | BioLegend |
| CD45.1 | Alexa Fluor® 700 | A20 | 110724 | BioLegend |
| CD45.2 | PE/Cyanine7 | 104 | 109830 | BioLegend |
| CD49a | Brilliant Violet 510 | Ha31/8 | 740144 | BD Biosciences |
| CD69 | FITC | H1.2F3 | 104506 | BioLegend |
| CD8a | Brilliant Violet 711 | 53-6.7 | 100759 | BioLegend |
| CD8a | Biotin | 53-6.7 | 553028 | BD Biosciences |
| B220 | Brilliant Violet 785 | RA3-6B2 | 103246 | BioLegend |
| B220 | Biotin | RA3-6B2 | 103204 | BioLegend |
| Blimp1 | Alexa Fluor 647 | 5E7 | 150004 | BioLegend |
| CXCR5 | Biotin | SPRCL5 | 13-7185-82 | Thermo Fisher Scientific |
| F4/80 | FITC | BM8 | 123108 | BioLegend |
| GL-7 | PE/Cyanine7 | GL7 | 144620 | BioLegend |
| Human IgG (H+L) | Alexa Fluor 647 | N/A | 109-606-088 | Jackson ImmunoResearch |
| IFN-γ | Brilliant Violet 605 | XMG1.2 | 505840 | BioLegend |
| IFN-γ | PE | XMG1.2 | SKU 50-7311 | Cytek Biosciences |
| IgA | FITC | Polyclonal | 1040-02 | Southern Biotech |
| IgD | Brilliant Violet 711 | 11-26c.2a | 405731 | BioLegend |
| IgG | Brilliant Violet 421 | Poly4053 | 405317 | BioLegend |
| IgG | Brilliant Violet 510 | Poly4053 | 405331 | BioLegend |
| IgM | APC-eFluor™ 780 | II/41 | 47-5790-82 | Thermo Fisher Scientific |
| IL-4 | PE/Cyanine7 | 11B11 | 25-7041-82 | Thermo Fisher Scientific |
| IL-17a | Alexa Fluor 700 | TC11-18H10.1 | 506914 | BioLegend |
| IL-21 | IL-21R Fc Chimera | N/A | 596-MR | R&D Systems |
| Ly6C | Brilliant Violet 650 | HK1.4 | 128049 | BioLegend |
| Ly6G | Brilliant Violet 711 | 1A8 | 127643 | BioLegend |
| PD-1 | Brilliant Violet 711 | 29F.1A12 | 135231 | BioLegend |
| RBD | PE | N/A | N/A | House-made |
| RBD | APC | N/A | N/A | House-made |
| Siglec-F | Brilliant Violet 421 | S17007L | 155509 | BioLegend |
| Spike<sub>539-546</sub> MHC I tetramer | PE | N/A | N/A | NIH Tetramer Core |
| Spike<sub>539-546</sub> MHC I tetramer | APC | N/A | N/A | NIH Tetramer Core |
| Streptavidin | Brilliant Violet 421 | N/A | 405225 | BioLegend |
| TER-119 | Biotin | TER-119 | 553672 | BD Biosciences |
| TGF-beta RII | PE | N/A | FAB532P | R&D Systems |
| TNF-α | PE/Cyanine7 | MP6-XT22 | 506324 | BioLegend |
| Viability | Zombie Aqua | N/A | 423102 | BioLegend |
| Viability | Zombie NIR | N/A | 423106 | BioLegend |

